# Candida-Klebsiella interactions rewire fungal morphogenesis to create a host environment favoring pathogen survival with enhanced tissue pathology

**DOI:** 10.64898/2026.05.06.723213

**Authors:** Tamires Bitencourt, Irina Tsymala, Rafael de Freitas Silva, Nicholas Wedige, Celine Prakash, Trinh Phan-Canh, Sabrina Villar Pazos, Filomena Nogueira, Sandra Högler, Theodora Ziu, Philipp Penninger, Debora Coraça-Huber, Michaela Lackner, Wilfried Ellmeier, Thomas Lion, Karl Kuchler

## Abstract

Clinically relevant infections commonly develop within polymicrobial environments where interkingdom interactions shape host responses and disease trajectories. *Candida albicans* and *Klebsiella pneumoniae* are critical pathogens that can co-exist in the respiratory tract, yet the consequences of their interaction in terms of fungal physiology, pathogenicity and impact on disease outcomes remain poorly understood. Here, we show that *K. pneumoniae* enhances *C. albicans* virulence traits suggesting that co-infections could exacerbate lung disease. Mechanistically, bacterial presence induces fungal hyphal morphogenesis via MAPK–CEK signaling, coupled to metabolic rewiring and alterations in cell wall remodeling and septation, resulting in highly elongated hyphae that escape faster from macrophages. At the host level, co-infection reprograms macrophages into a non-canonical state characterized by overlapping pro- and anti-inflammatory modules, integrating type I interferon and IL-10 signaling. This response contributes to tissue damage and facilitates fungal persistence. Our findings reveal that the bacterial–fungal interactions coordinately reprogram pathogen behavior and host immunity, promoting pathogenic synergy and potentially conferring a negative impact on disease outcomes.

**HIGHLIGHTS:** *Candida–Klebsiella* interactions modulate hyphal morphogenesis.

Ectopic morphogenesis encompasses septation, cell wall remodeling and carbon metabolism.

*Candida–Klebsiella* co-infections trigger tissue hyperinflammation and compensatory regulation.

*Candida-Klebsiella* co-infection establishes a host environment facilitating microbial dissemination and tissue pathology.

*Candida–Klebsiella* interactions enhance fungal virulence potentially impacting the severity of co-infections

## INTRODUCTION

*Candida albicans* is a pleiomorphic human commensal and normal part of the microbiota of immunocompetent individuals, colonizing mucosal tissues particularly in the respiratory and gastrointestinal tracts^1^. However, it can also cause diseases ranging from superficial mucosal infections to life-threatening invasive candidiasis in immunocompromised patients, accounting for an estimated 400,000 cases annually worldwide^2^. The transition from commensalism to pathogenicity is driven by complex multifactorial determinants, including host immune defense, as well as interactions with resident microbiota^3–5^. These interactions remodel local environmental cues, thereby influencing *C. albicans* morphogenesis, metabolism and virulence traits.^6–9^ Defining how bacterial communities potentially constrain or promote fungal pathogenicity is therefore central to understanding susceptibility to invasive candidiasis.

Infections often develop within complex polymicrobial environments, mostly facilitated by biofilm formation, where interspecies interactions shape pathogen behavior and host responses^10^. There is growing recognition that polymicrobial infections represent a major threat, owing to complicated clinical diagnosis, challenging antimicrobial treatments and variable outcomes^10–12^. Despite these insights, the mechanisms through which bacterial–fungal interactions regulate fungal fitness, immune evasion and host defense remain poorly defined. Recent studies demonstrate that commensal bacteria can induce marked changes in the *C. albicans* cell wall^13^, thereby altering immune responses. Further, fungal-bacterial co-culture modulates hyphal morphogenesis during macrophage interactions^8^. However, whether such interspecies interactions rewire fungal transcriptional programs, alter host immune responses and influence infection outcome *in vivo* remains unresolved.

Host cells discriminate between commensal and pathogenic states of *C. albicans*, whereby invasive infections are commonly initiated by hyphal morphogenesis and penetration of epithelial and mucosal barriers. Dissemination triggers local innate immune responses, including recruitment of phagocytes to infected tissues, with neutrophils and macrophages playing a central role in fungal control^14^. Macrophages deploy multiple antifungal strategies, including phagocytosis, nutrient restriction, cytokine production, and the generation of reactive oxygen and nitrogen species^15^. Further, they can physically constrain elongating hyphae at septal junctions to limit fungal escape^16^. Neutrophils likewise contribute critically to fungal control by killing hyphal forms through NET (neutrophil extracellular traps) formation and myeloperoxidase-dependent mechanisms^17^. In response, *C. albicans* can activate immune evasion countermeasures promoting survival within the phagosome. For example, metabolic rewiring can support growth under nutrient limitation, and remodeling of the fungal cell wall counteracts fungal killing^18,19^. Additionally, intracellular hyphal elongation can lead to phagosomal rupture and macrophage lysis, facilitating fungal escape and dissemination^20^. Because the same myeloid populations also coordinate antibacterial defense, bacterial-fungal co-infection may reprogram these responses to either exacerbate inflammatory tissue injury or impair fungal clearance. Defining such context-dependent host responses in clinically relevant infection models is therefore essential for understanding polymicrobial disease and for informing therapeutic strategies.

In previous work, we investigated interactions between *Aspergillus fumigatus* and *Klebsiella pneumoniae (K. pneumoniae)* and demonstrated a remarkable bacterial-driven rewiring of fungal carbon metabolism, raising important questions about how metabolic reprogramming may impact the host response^21^. *K. pneumoniae* is part of the ESKAPE (*Enterococcus faecium*, *Staphylococcus aureus*, *Klebsiella pneumoniae*, *Acinetobacter baumannii*, *Pseudomonas aeruginosa* and other members of the *Enterobacterales spp*) group of organisms and a major cause of hospital-acquired infections.^22,23^ The World Health Organization (WHO) defines it as a priority bacterial pathogen^24^. *K. pneumoniae* commonly colonizes mucosal surfaces, including the respiratory tract, making interactions with *C. albicans* in the lungs a clinically intriguing, yet underexplored scenario.

Here, we investigate how *K. pneumoniae* reshapes *C. albicans* physiology, virulence-associated traits and host responses during co-infection and address the potential implications of this interaction for affecting the severity of disease and outcome. We show that bacterial-fungal co-cultures drive profound remodeling of *C. albicans* biofilms, as indicated by altered hyphal morphogenesis, impaired septation, cell-cycle alterations, and changes in the fungal mannan cell wall distribution. These changes promote thicker biofilms with increased capacity for dispersal. Moreover, interspecies interactions generate a permissive host immune environment, in which the macrophage activation state is characterized by overlapping pro- and anti-inflammatory programs that may favor fungal survival and immune evasion. *In vivo*, co-infections in an acute disease model drive severe pulmonary immunopathology, accompanied by extensive fungal hyperfilamentation and enhanced bacterial dissemination. Together, our findings identify bacterial-fungal interactions as key determinants of fungal pathophysiology and host response and highlight the need to consider polymicrobial contexts when studying and treating invasive infections.

## RESULTS

### *C. albicans* reprograms growth and carbon metabolism in response to *K. pneumoniae*

To investigate the dynamics of interspecies interactions between *C. albicans* and *K. pneumoniae*, we established a mixed biofilm model in which both microbes were co-cultivated for 24 h prior to analysis. Interactions were evaluated under two conditions, RPMI 1640, a mammalian medium that mimics physiological conditions and supports *C. albicans* growth and biofilm formation^25^, and M9 minimal medium, enabling assessment of microbial interactions under resource restriction, including nutrient competition and cross feeding.

Biofilms in both conditions were then subjected to dual-species RNA-seq^26,27^, and transcriptional dynamics were assessed with a focus on the *C. albicans* response (Fig. 1a). Notably, interspecies effects were more pronounced in RPMI, as reflected by clearer separation in the PCA (Fig. 1b; Extended Fig. 1a), indicating enhanced transcriptional responsiveness and adaptive shifts to bacterial exposure under this condition.

**Figure 1.**
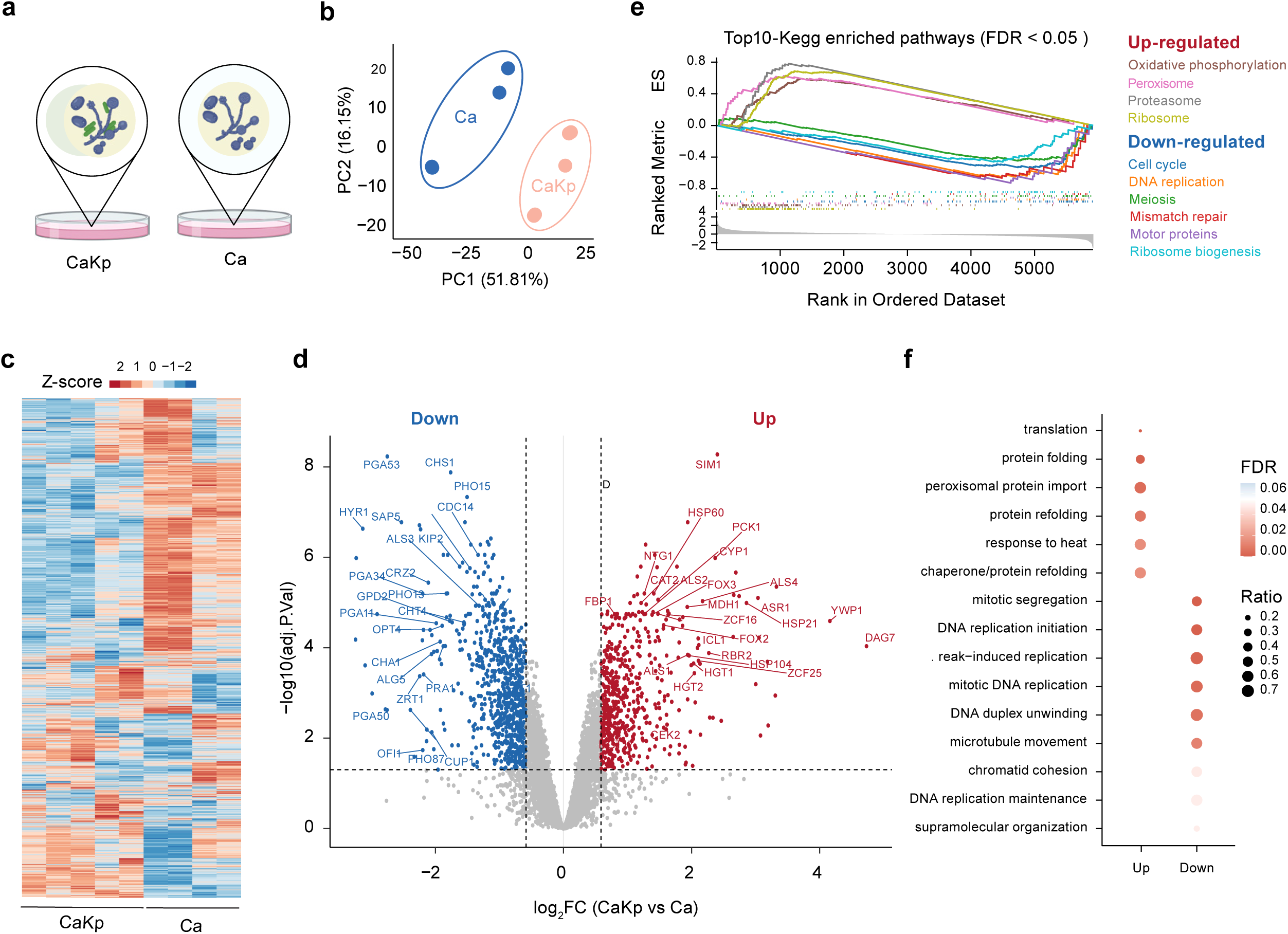
Transcriptional dynamics of *C. albicans* in mixed biofilms with *K. pneumoniae* cultured in RPMI medium. **(a)** Biofilm growth conditions submitted to dual RNA-seq analysis comparing *C. albicans* alone (Ca) *versus Candida-Klebsiella* (Cakp), after growing in biofilm mode for 24h. **(b)** Principal component analysis (PCA) of transcriptomes from mixed and single pathogen cultures of *C. albicans* (Ca) and *C. albicans* with *K. pneumonia* (CaKp). **(c)** Euclidean hierarchical clustering of gene expression levels across all genes following Trimmed Mean of M-values (TMM) normalization. Expression values are shown as log-counts per million (logCPM) and were scaled by Z-score. Data from least four independent biological replicates are shown for single Ca cultures and CaKp co-cultures**. (d)** Volcano plot representing up- and downregulated differentially expressed genes (DEGs) from the comparison CaKp *vs* Ca, using log2-fold change (log2FC) *versu*s −log10 adjusted P value for all genes. DEGs were defined using log2FC±0.5 cut-off. Top selected genes are shown. **(e)** Gene Set Enrichment Analysis (GSEA) covering the KEGG pathways (https://www.kegg.jp/kegg-bin/show_organism?org=cal) for *C. albicans* (CaKp *vs* Ca). Pathways with adjusted P value (Benjamini–Hochberg FDR) < 0.05 were considered as significantly enriched. **(f)** Functional enrichment analysis related to up- and down-regulated biological processes based on Gene ontology (GO) terms of DEGs in CaKp compared to Ca alone.

*C. albicans* exhibited a robust transcriptional response to the presence of *K. pneumoniae* in RPMI, with 1,360 differentially expressed genes (DEGs), including 584 upregulated and 776 downregulated genes (Figs. 1c-d). Analysis of the modulated gene sets, together with gene set enrichment analysis (GSEA) and Gene Ontology (GO) analysis, indicates that this response reflects a coordinated adaptive program involving metabolic reprogramming, induction of heat shock proteins and protein-folding stress responses, and attenuation of cell cycle–associated processes (Figs. 1e-f).

Among the highly upregulated genes in *C. albicans* in response to *K. pneumoniae* were those associated with cell wall and cell surface functions (*YWP1, DAG7, ALS4*), morphogenesis-related signaling pathways (*CEK2, ZCF25*), alternative carbon metabolism (*PCK1, FBP1, ICL1, FOX3, CAT2*), and heat shock response proteins (Fig. 1d). In contrast, downregulated genes included factors involved in cell surface remodeling, nutrient acquisition, and virulence-related functions, including *HYR1, PRA1, ZRT1, CRZ2, CHS1, CHT4,* and *PGA53*.

In agreement with these findings, the top 100 Z-score–ranked genes reinforced this adaptive transcriptional program, highlighting the activation of morphogenesis, stress response, and metabolic reprogramming. Conversely, genes associated with cell cycle progression, cell wall organization, and metal ion uptake were reduced, indicating a shift away from proliferation toward a survival- and morphogenesis-associated state compatible with phenotypic plasticity (Supplementary Fig. S1a).

In contrast, *C. albicans* biofilms grown in M9 responded to *K. pneumoniae* by exhibiting a transcriptional profile skewed toward ion acquisition and metal homeostasis, indicative of increased demand for essential nutrients. This is also consistent with a more constrained state that may delay morphogenetic progression. Upon co-culture with *K. pneumoniae*, *C. albicans* displayed 1,250 DEGs, including 644 upregulated and 606 downregulated genes (Extended Figs. 1b-c). Upregulated genes included those involved in alternative carbon metabolism (e.g., *PCK1, FBP1, CIT1,* and *ACS1*) as well as nutrient uptake and environmental adaptation (e.g., *TRY4–TRY6, PRA1, ZRT1, ZRT2, PHO84, PHO86, HGT16,* and *CSA2*). In contrast, downregulated genes were enriched for cell wall–associated functions, including *PGA11, PGA23, PGA30, PGA52, PGA53, MNN1, XOG1, CHS1, CHS2,* and *SAP7* (Extended Fig. 1c; Fig. S1a). Consistent with these patterns, pathway enrichment analysis revealed activation of iron acquisition, macromolecule catabolism, RNA processing, the glyoxylate cycle, and peroxisomal functions, alongside reduced expression of genes associated with glucose utilization and cell cycle–related processes (Extended Figs. 1d-e).

Although the primary focus of this study was the *C. albicans* response to *K. pneumoniae*, we also examined the bacterial transcriptional landscape to better contextualize a potential impact on fungal behavior. Clustering analysis revealed a clear separation between *K. pneumoniae* monocultures and mixed-species biofilms, indicating a substantial transcriptional reprogramming in response to the presence of *C. albicans* across both media conditions (Fig. S1b). In RPMI, *K. pneumoniae* exhibited enrichment of pathways associated with phosphotransferase system (PTS) activity and ribosomal function, alongside with a marked downregulation of amino acid metabolism genes (Figure S1c). Specifically, genes involved in tryptophan, histidine, and branched-chain amino acid synthesis (*trpA-F, hisD, alsS*) were repressed, whereas nucleotide metabolism and translation factors (*deoA, deoC, rpmJ, rpmH*) were upregulated (Fig. S1d). These bacterial transcriptional adjustments likely can alter the local microenvironment by affecting availability of certain amino acids and nucleoside metabolites, while sustaining active growth and protein synthesis.

Likewise, in M9, the *K. pneumoniae* response entailed activation of central carbon metabolism, including glycolysis and gluconeogenesis, together with increased ribosomal activity. This was accompanied by downregulation of quorum sensing and aromatic compound metabolism (Fig. S1c). Upregulated genes included *mrk operon*, phosphate uptake (*pst operon*), ion transport (*kdp operon*), and translation (*rpl genes*), consistent with a need for enhanced nutrient acquisition to maintain growth. Meanwhile, genes linked to alternative nutrient pathways and transport systems (*rut, ace, mal, mgl*) were repressed (Fig. S1d). This data suggest that bacteria sustain robust growth and restrict resource availability for *C. albicans*, effectively constraining fungal development.

### Bacterial–fungal interactions reshape transcription and biofilm architecture

Given that changes in carbon metabolism and morphogenetic programs represents a marked strategy in *Candida* responses to *Klebsiella*, and that both are linked to cell wall remodeling, we next examined transcriptional changes in cell wall–associated pathways, including filamentation, cell cycle progression, cell wall integrity (CWI) mediated by MAPK signaling, and carbon metabolism. These processes are coordinately regulated during biofilm development^28–30^. This analysis was performed by comparing mixed *C. albicans–K. pneumoniae* (CaKp) and single-species *C. albicans* (Ca) biofilms in RPMI (Fig. 2a).

**Figure 2.**
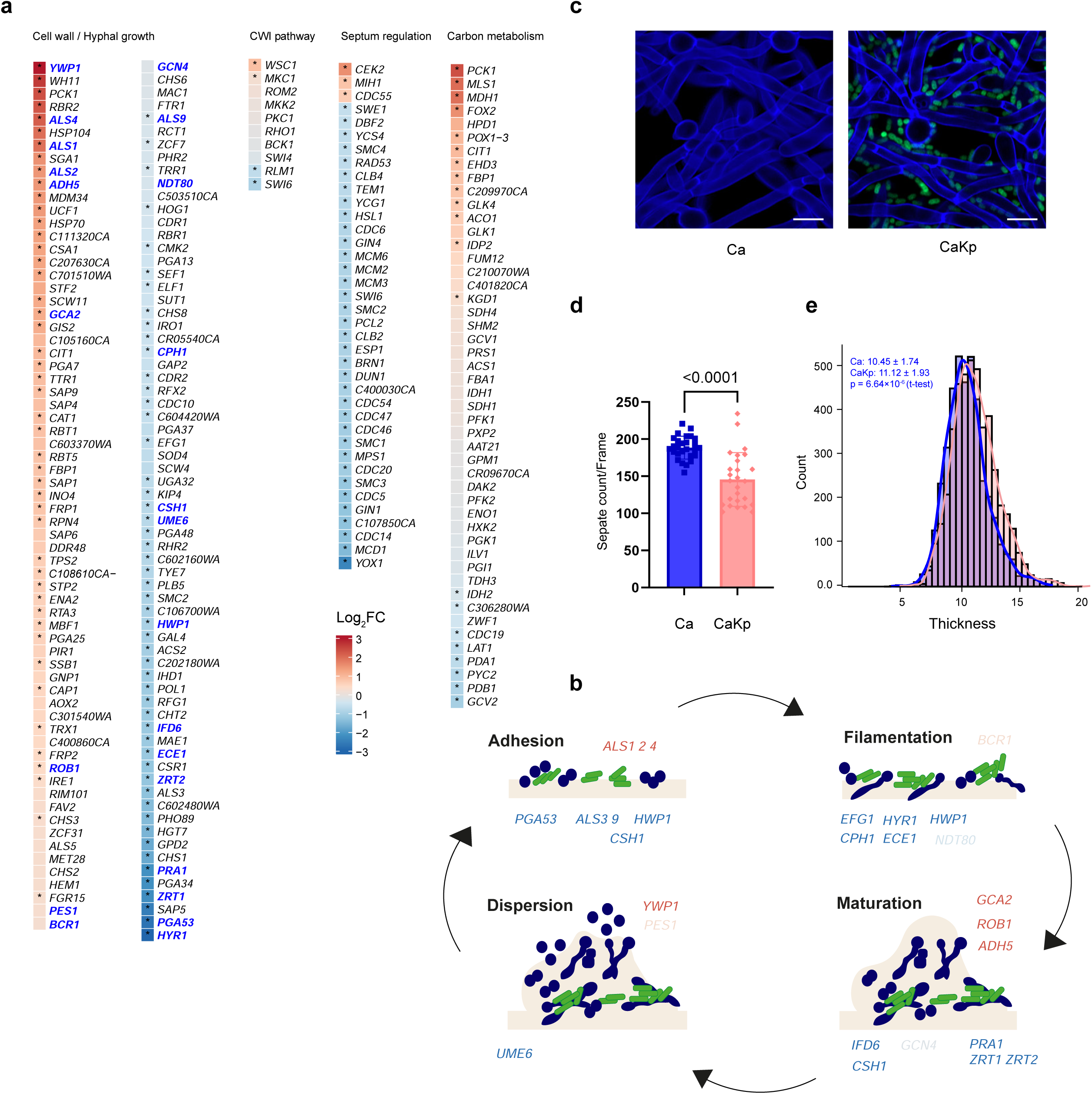
Pathway-resolved transcriptional changes induced by bacterial–fungal interactions that impact Candida biofilm formation, architecture, and morphogenesis programs in RPMI. **(a)** DEGs of selected categories such as hyphal growth/cell wall, CWI pathway, cell cycle and carbon metabolism in CaKp *vs* Ca conditions. Color scales indicate log2FC. **(b)** Schematic representation of biofilm stages, and DEGs potentially involved in each stage following interaction with *Klebsiella*. Selected genes are highlighted in blue in the cell wall and hyphal growth heat maps. **(c)** Biofilms of Ca and Cakp were stained with Calcofluor White (CFW 25 µg/mL) and co-cultured with a GFP-tagged *Klebsiella* strain. Biofilms were analyzed after 1.5 h of adhesion followed by 6 h of growth. **(d)** Septa counts were quantified using fluorescence images of mixed biofilms and compared with single-species Ca biofilms. **(e)** Biofilm thickness was measured per z-stack using 100 randomly selected stacks (400 × 400 px). Data were collected from three biological replicates per condition, with batch sizes ranging from 10 to 48. Statistical significance using an unpaired t-test; **** indicates p < 0.0001.

*C. albicans* responded to *K. pneumoniae* with coordinated upregulation of genes associated with adhesion, stress adaptation, proteolysis, and cell wall remodeling. These included adhesins (*ALS1, ALS2, ALS4*), stress response factors (*HSP70, HSP104, CAT1, TRX1, IRE1*), proteases (*SAP1, SAP9*), cell wall–associated genes (*CHS3, PGA7, PGA25*), and the key biofilm regulator *ROB1*. In contrast, central regulators of hyphal development and septation (*HYR1, ECE1, HWP1, EFG1, UME6, CHS1*) as well as zinc acquisition genes (*PRA1, ZRT1, ZRT2*) were downregulated (Fig. 2a).

Genes involved in cell cycle regulation were also predominantly repressed, with the exception of *MHI*, *CDC5*, and the CEK2 MAPK pathway gene, which were significantly upregulated. Within the cell wall integrity (CWI) signaling pathway, the sensor *WSC1* and the MAP kinase *MKC1* were induced, whereas the transcriptional co-regulators *SWI6* and *RLM1*, which contribute to cell cycle progression and septation, were downregulated (Fig. 2a). Together, these changes indicate a rewiring of morphogenesis programs, likely coupled to activation of cell wall remodeling and damage-response signaling programs^31^.

Mixed biofilms also displayed a pronounced shift in overall carbon metabolism, a feature that appeared as a hallmark of bacterial-fungal interactions^13^, including in our previous report on the interaction between *A. fumigatus* and *K. pneumoniae*^21^. Indeed, we observed increased expression of genes involved in β-oxidation, glyoxylate cycle, and gluconeogenesis, but reduced expression of genes linking pyruvate and amino acid metabolism to mitochondrial carbon flux (Fig. 2a).

Biofilm formation in *C. albicans* progresses through four main stages (adhesion, filamentation, maturation and dispersion)^32^. To assess how *K. pneumoniae* influences biofilm formation in *C. albicans* we mapped differentially expressed genes onto distinct biofilm stages-related steps. After 24h of interspecies interaction, CaKp biofilms showed a repression of adhesion- and filamentation-associated processes, accompanied by partial induction of matrix- and dispersion-related steps (Fig. 2b). Indeed, our data highlighted the expression and regulation of a set of dispersion genes, such as *PES1, UME6,* and *YWP1*^33^. Overall, these data suggest that the presence of *K. pneumonia* alters the temporal biofilm-associated transcriptional program of *C.albicans*, with repression of early-stage genes and partial enrichment of later-stage programs (Fig. 2b).

To further assess the impact of nutrient availability on *Candida* mixed-biofilm dynamics, we extended the analysis to M9 medium and compared pathway responsiveness between single- and mixed-species biofilms. Notably, these data suggest that under nutrient-limited conditions, *C. albicans* maintains a metabolically active state while attenuating cell cycle progression. In this context, cell cycle regulation appeared tightly controlled rather than fully arrested, as reflected by the upregulation of transcriptional regulators such as *SWI4* and *SWI6* (Extended Fig. 2a), accompanied with the upregulation of a distinct set of filamentation- and zinc acquisition–associated genes, including key hyphal regulators such as *ECE1, HYR1, TUP1,* and *TEC1* (Extended Fig. 2a). Mapping these regulated genes onto biofilm developmental programs revealed a transcriptional pattern consistent with delayed progression through biofilm development, with programs associated with the filamentation stage being particularly enriched (Extended Fig. 2b).

We next performed fluorescence-based imaging of single and mixed biofilms grown in RPMI to assess the impact of *K. pneumonia* on *C. albicans* biofilm architecture (Fig. 2c). Mixed CaKp biofilms exhibited significantly reduced septation when compared to single-species Ca biofilms (Fig. 2d). This is consistent with the observed changes in cell cycle regulation and hyphal development. Despite reduced septation, mixed biofilms displayed increased thickness at 6 h post-adhesion (Fig. 2e), suggesting enhanced hyphal reorganization in the presence of *K. pneumoniae*, consistent with altered biofilm formation kinetics.

Given the established role of lipids and sterol-rich membrane domains in hyphae transition and biofilm maturation^34,35^, we next examined ergosterol metabolism in mixed biofilms. Genes involved in ergosterol biosynthesis (*ERG7, ERG6, ERG24, ERG 25, ERG5, ERG4*) were broadly repressed in CaKp biofilms (Supplementary Fig. 2a). Consistent with this transcriptional profile, filipin III staining revealed altered ergosterol distribution pattern in mixed biofilms, showing a preferential apical localization although no difference in overall fluorescence intensity were observed. Of note, upon exposure to amphotericin B, an ergosterol-binding antifungal, filipin fluorescence increased in single-species biofilms but decreased in mixed biofilms, suggesting differential sterol accessibility and membrane organization (Supplementary Fig. 2b).

Moreover, mannans represent a major complex carbohydrate of the *C. albicans* biofilm matrix^35,36^, and since it can be differentially regulated by bacteria ^13^. We therefore assessed the distribution of cell wall mannans in mixed biofilms using concanavalin A (ConA) staining. Compared with Candida biofilms, CaKp biofilms exhibited increased ConA fluorescence (Supplementary Fig. 2c). This suggests, that *Klebsiella* may alter fungal cell wall or matrix properties, with potential consequences for host immune recognition.

### Co-infections with *C. albicans* and *K. pneumoniae* alter host immunity and disease

*K. pneumonia* modulates *C. albicans* transcriptional programs *in vitro* particularly those linked to morphogenesis and cell wall remodeling. This may enhance fungal fitness and pathogenic potential within host tissues. To determine how this interaction shapes infection outcome *in vivo*, we established a model of acute fungal pneumonia in C57BL/6J mice by intranasal instillation. We compared disease progression and outcome across different infection scenarios: *C. albicans* infection only, simultaneous co-infection with CaKp, and secondary *C. albicans* infection following *K pneumoniae* (Kp-Ca) (Fig. 3a).

**Figure 3.**
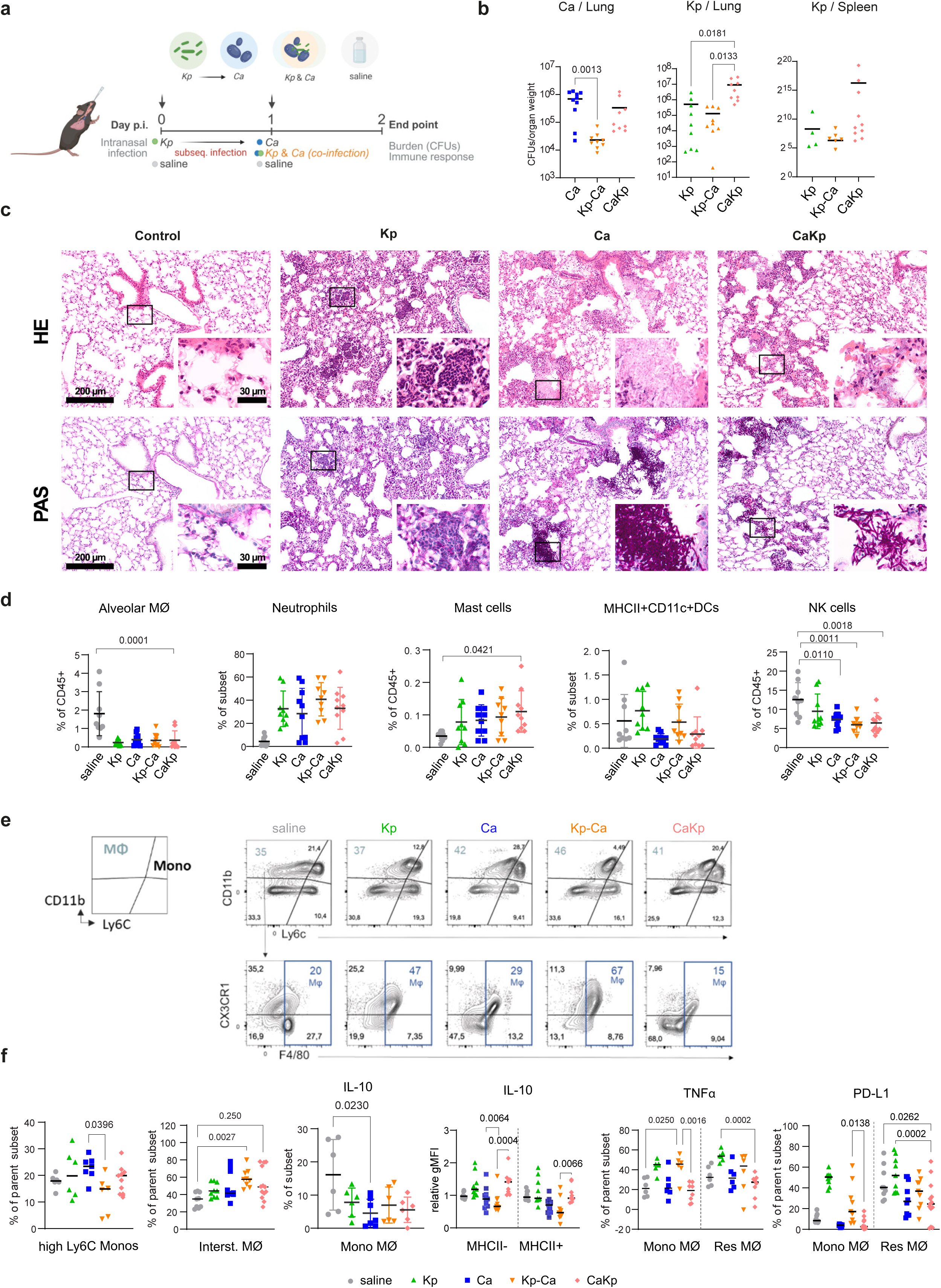
Microbial co-infection exacerbates lung damage, favors *Klebsiella* dissemination and modulates macrophage recruitment. **(a)** Schematic representation of the murine nasal infection model used. **(b)** Fungal burden 2 days p.i., determined by CFUs quantification of lungs/spleen from infected animals normalized to organ weight. Each dot represents one animal pooled from 3 independent experiments. Mean ± SD is shown. **(c)** Inflammation and immune cell recruitment are shown by H&E staining of lung sections at 2 days p.i. from two animals per group; periodic acid-Schiff (PAS) staining was performed for visualization of fungal cells and the co-localization of Ca and Kp. Black dotted rectangles indicate areas chosen for magnification. **(d)** Innate immune cell recruitment profiled by flow cytometry following *ex vivo* stimulation of lungs cells with heat-killed *Candida-Klebsiella* at 2days p.i. **(e)** Gating strategy to identify CD11b⁺ and Ly6C⁺ subpopulations in pooled lung immune cells. **(f)** Proportion of monocytes and macrophages producing IL-10 or TNF-α, and expressing PD-L1, in mouse lungs after coinfection.

Mouse body weight loss was comparable between single (*C. albicans* alone) and co-infected (CaKp) groups, as well as between *K. pneumonia* (Kp) and Kp-primed *C. albicans* (Kp-Ca) groups. However, when directly compared to the CaKp group, Kp mice exhibited the most pronounced weight loss already at 1 day post-infection (p.i.) (Supplementary Fig. 3a). The quantification of fungal burdens revealed a reduced *C. albicans* abundance in lungs of Kp-Ca mice, with a similarly decreasing trend observed in CaKp mice (Fig. 3b). In contrast, bacterial loads were significantly elevated in CaKp mice, accompanied by increased dissemination into the bloodstream, suggesting more severe tissue damage and loss of barrier integrity (Fig. 3b). Consistent with that, CaKp mice developed increased lung and spleen weights suggesting enchanced inflammatory response and active immune defence (Fig. S3b). To collect histopathological evidence, we also analyzed lung sections from all infection conditions (Fig. 3c). Control animals showed no lesions in the lungs. PAS staining revealed fungal infiltrates in all *Ca-*infected groups, with PAS-positive bacterial colonies frequently co-localizing with fungal hyphae in co-infected mice. Inflammatory foci were predominantly peribronchiolar but extended deeply into the alveolar parenchyma in CaKp animals, where barrier architecture was disrupted by fungal invasion (Fig. 3c). Compared to single infection or Kp-Ca, or even the Kp-CaKp group (Supplementary Fig. 3c), CaKp lungs exhibited larger hyphal clusters extending into the parenchyma. Collectively, these results showed that co-infection promotes invasive disease with substantial tissue destruction. However, priming animals with *K. pneumoniae* challenge protected against subsequent *C. albicans* infection.

Histology images also revealed the infiltration of neutrophils and macrophages by Kp, while Kp and Ca showed pronounced lymphocyte infiltrates, which were less evident by co-culture of CaKp (Fig 3c). To better profile the immune cell recruitment and response to the different infection conditions, we next used spectral flow cytometry (Figs 3d, 3e and 3f). Across all infected groups, we observed a reduction in the frequency of alveolar macrophages, consistent with cell loss or their phenotypic M1-M2 conversion during acute inflammation. Neutrophil recruitment was robust across all conditions, reflecting intact antimicrobial defence. Mast cells showed a modest increase in co-infected mice, while dendritic cells (DCs) and natural killer (NK) cells were largely unchanged across fungal infection conditions (Figs 3d, 3e and 3f). Notably, Ly6C^high^ inflammatory monocytes were not prominently induced in Kp-challenged mice (Figure 3e). However, both co-infection conditions (CaKp and Kp-Ca) displayed an expansion of CX3CR1^+^ interstitial macrophages, implying monocyte-to-macrophage differentiation. Ex *vivo* restimulation of lung leukocytes with heat-inactivated pathogens revealed enhanced anti-inflammatory IL-10 production in co-infected mice, which was largely absent in the Kp-Ca group (Fig. 3f). TNFα expression was also elevated, particularly in Ly6C^+^ monocyte-derived macrophages. Although *Kp* infection may upregulate PD-L1 expression as a mechanism of immune suppression^37^, we observed that *Candida* infection in fact reduced PD-L1^+^ macrophage subsets. Together, these findings showed that *K. pneumoniae* infection modulates immune cell recruitment and macrophage polarization, revealing exacerbated inflammation and tissue damage during co-infection. These effects appeared to be at least partially driven by fungal adaptation and altered immune cell-microbe interactions in the lung tissue microenvironment.

### Combined *Candida–Klebsiella* infection triggers regulated macrophage responses

Given that co-infection triggered extensive tissue damage *in vivo*, we next investigated these interactions in a controlled setting using bone marrow–derived macrophages (BMDMs) to assess how fungal and bacterial pathogens collectively influence macrophage responses. BMDMs were infected with each pathogen individually or in combination for 2 h, and then cells were harvested for bulk RNA-seq analysis. Comparison of gene expression following *Candida* or *Candida–Klebsiella* co-cultures revealed a broad set of upregulated genes, with relatively few downregulated (Figs. 4a-b). Notably, CaKp exposure elicited an even stronger response, characterized by marked upregulation of chemokines (*Ccrl2*, *Ccl3*, *Ccl4*, *Ccl2*, *Cxcl2*, *Cxcl1*, *Cxcl10*, and *Ccl7*), pro-inflammatory cytokines (*Il12*, *Il6*, *Lif*, *Il1a*, *Il1b*, *Ptgs2*, *Tnf*), immune regulators (*Il10*, *Tnfaip3*, *Acod1*, and *Il1rn*), NF-kB pathway (*Nfkbia*), survival factor (*Bcl2a1a*) and leucocyte recruitment (*Icam1*) (Figs 4a–b). Similarly, *Klebsiella* infection alone also triggered a robust transcriptional host response (Supplementary Fig 4a).

**Figure 4.**
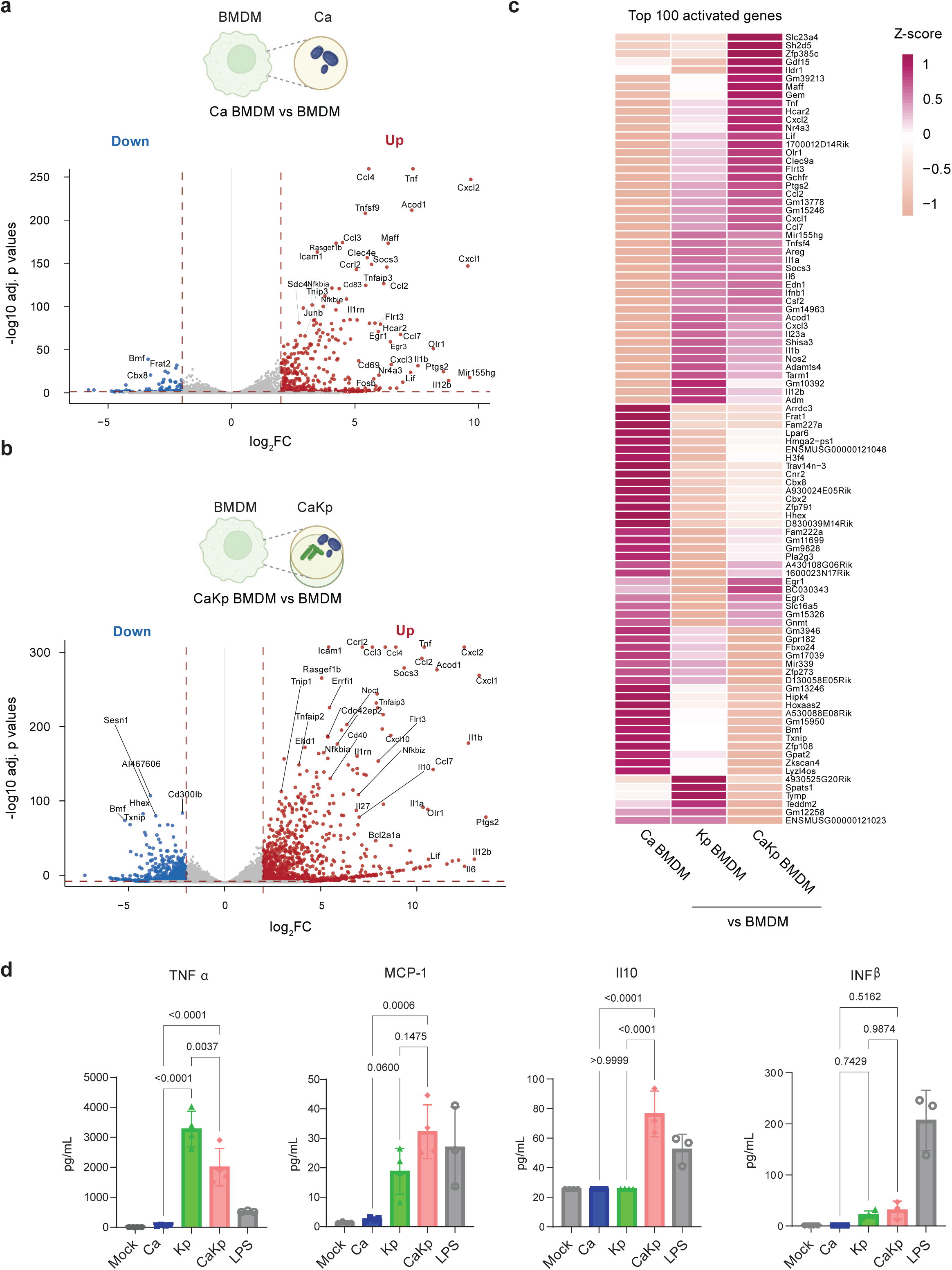
Macrophage response to microbial co-cultures triggers a balance between pro-inflammatory and anti-inflammatory programs. **(a)** Volcano plot representing the up- and down-regulated DEGs from CaBMDM *vs* BMDM or CaKpBMDM *vs* BMDM **(b)**; log2-fold change (log2FC) *versu*s −log10-adjusted P value for all genes. **(c)** Top 100 differentially responsive genes across CaBMDM, KpBMDM, and CaKpBMDM conditions compared to BMDMs alone, with Z-score–scaled expression values. **(d)** Cytokine secretion by BMDMs following 2 h challenge with *K. pneumoniae* (Kp), *C. albicans* (Ca), or co-culture (CaKp) using a multiplicity of infection (MOI) of 3:1.

GSEA of host transcriptional responses revealed that *Candida* exposure predominantly enriched inflammatory processes related to interleukin-1 responses and granulocyte migration, whereas combined *Candida–Klebsiella* exposure was associated with enrichment of chemokine-mediated signaling pathways (Supplementary Figs 4b-c). In contrast, *Klebsiella* infection alone primarily triggered processes related to granulocyte chemotaxis and migration (Supplementary Fig 4d).

Notably, CaKp exposure further amplified the expression of pro-inflammatory mediators, including *Ptgs2, Tnf, Cxcl1,* and *Ccl2*, along with regulatory and interferon-responsive genes such as *Il1rn, Rsad2,* and *Il10*. Consistently, analysis of the top 100 most highly modulated transcripts identified activation of specific subset of genes (Fig 4c), highlighting the activation of the following genes mainly for CaKp group (*Ccl7*, *Cxcl1*, *Ccl2*, *Ptgs2*, *Flrl3*, *Clec9a*, *Lif*, *Cxcl2*, *Hcar2*, *Tnf*, *Maff*, *Ildr1*, *Gdf15*).

We further assessed cytokine secretion by BMDMs under the different infection conditions. Co-infection with CaKp induced markedly higher secretion of IL-10 and MCP-1 (CCL2), indicative of an anti-inflammatory and perhaps chemotactic response, whereas TNFα was elevated in both CaKp and *Klebsiella* single infections, with the highest levels observed in *Klebsiella* infection (Fig. 4d). Together, these data highlighted that CaKp exposure elicits a robust inflammatory response in macrophages that is actively counterbalanced by immunoregulatory pathways, resulting in tightly controlled macrophage activation states that facilitate *Candida* immune evasion.

### Co-infection promotes Candida hyphal escape with attenuated ROS response

To investigate how these effects on macrophage behavior affect the ultimate response to single *vs* mixed co-cultures exposure, and assess whether macrophage states do create a permissive environment for *Candida* escape, we determined *C. albicans* survival in BMDMs exposed to either *Candida* alone or *Candida–Klebsiella* (CaKp) (Fig. 5a). Notably, survival rates of *C. albicans* were significantly higher in BMDMs co-cultured with CaKp when compared to single-species exposure (Fig. 5b). We next examined phagocytosis rates. After 2 h of interaction, a considerable proportion of *Candida* from Ca group was internalized by macrophages, whereas a larger fraction of *Candida* cells from the CaKp group remained outside macrophages, suggesting either changes in the recognition patterns or potential hyphal escape (Figs 5c-d). Indeed, immunofluorescence and live-cell imaging revealed hyphal extensions beyond macrophage boundaries during CaKp exposure (Fig 5e), indicating that the presence of *K. pneumonia* triggers the elongated hyphal growth that facilitates evasion from phagocytosis (Supplementary Figs 5a-b).

**Figure 5.**
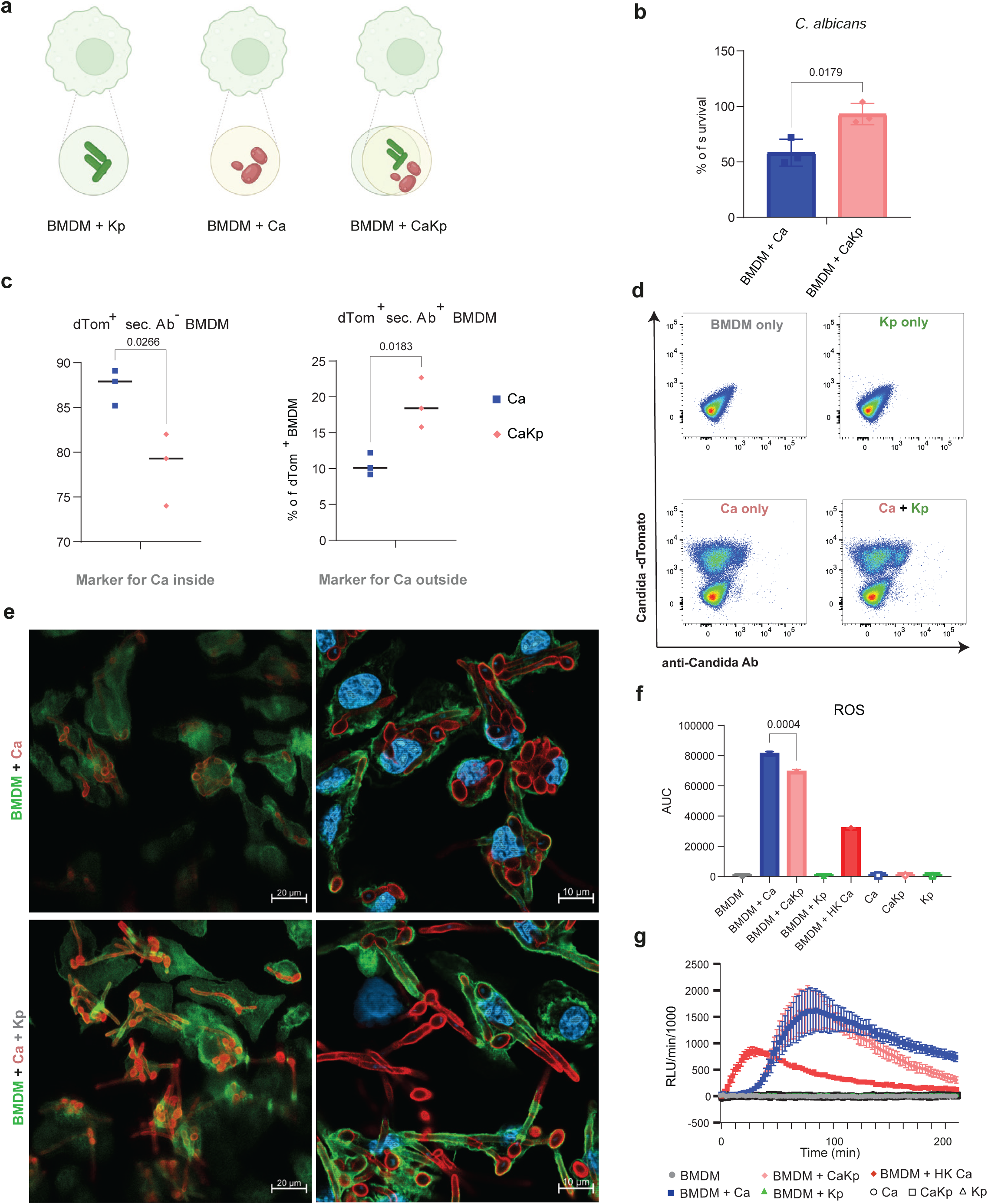
Klebsiella enhances *C. albicans* survival and hyphal escape with reduced macrophage ROS response. **(a)** Scheme of experimental conditions with BMDM exposed to *Klebsiella* (Kp), *Candida* (Ca) and co-infected with Ca and Kp (CaKp). **(b)** Fungal viability after 3h of co-culture with BMDMs shown as percentage values. Data represent the means and SD from three biological assays performed with four technical replicates. **(c)** Phagocytosis of dTom-labeled *C. albicans* by BMDMs after 2h of interaction assessed by flow cytometry. The means (horizontal lines) and SD from biological triplicates (dots) are shown. **(d)** Gating strategy for phagocytosis, to identify populations of Ca positive for dTOM and/or positive for anti-*Candida* Ab. **(e)** Immunofluorescence of BMDMs infected with *Candida* (Ca) or co-infected with Ca and *Klebsiella* (CaKp) for 2 h at an MOI of 2:1. Macrophages were stained with phalloidin, *Candida* with Texas Red–conjugated secondary antibodies, and nuclei with DAPI. (**f**) Area under the curve (AUC) for reactive oxygen species (ROS) production in BMDMs exposed to Ca, CaKp, Kp, heat-killed *C. albicans*, or pathogens alone as negative controls. (**g**) Real-time luminescence-based reactive oxygen species (ROS) assay with BMDMs co-cultured with Ca and CaKp, using heat-killed Ca as a control. Representative RLU of isoluminol over time per 1,000 BMDMs are depicted.

Considering that Klebsiella potentially induces changes in *C. albicans* morphogenesis and cell wall mannan distribution, and that reactive oxygen species (ROS) is a key macrophage defense against *C. albicans* hyphae, we quantified ROS production in BMDMs exposed to *Candida*, *Klebsiella*, or both, with heat-killed *Candida* serving as a positive control. *Klebsiella* alone did not trigger detectable ROS, whereas *Candida* induced strong and sustained ROS production. CaKp exposure elicited a weaker and less sustained ROS response in comparison to *Candida* alone (Figs f-g). Together, these results indicate that CaKp reduces *Candida* uptake, limits ROS-mediated killing, and promotes hyphal escape, while simultaneously skewing macrophages towards permissive states that allow for fungal immune evasion.

Collectively, these findings show that interaction with *K. pneumoniae* induces a distinct transcriptional and phenotypic state in *C. albicans*, marked by rewiring of yeast-to-hypha transition programs, carbon metabolism, stress responses, and cell wall-associated pathways. These changes led to altered fungal cell wall properties and enhanced hyphal extension, supporting a model in which bacterial co-infection promotes fungal adaptation and immune escape, as summarize in Fig 6. Our results highlight microbial interaction as an important determinant of fungal pathogenicity and host immune responses, with implications for understanding and potentially targeting polymicrobial infections.

**Figure 6.**
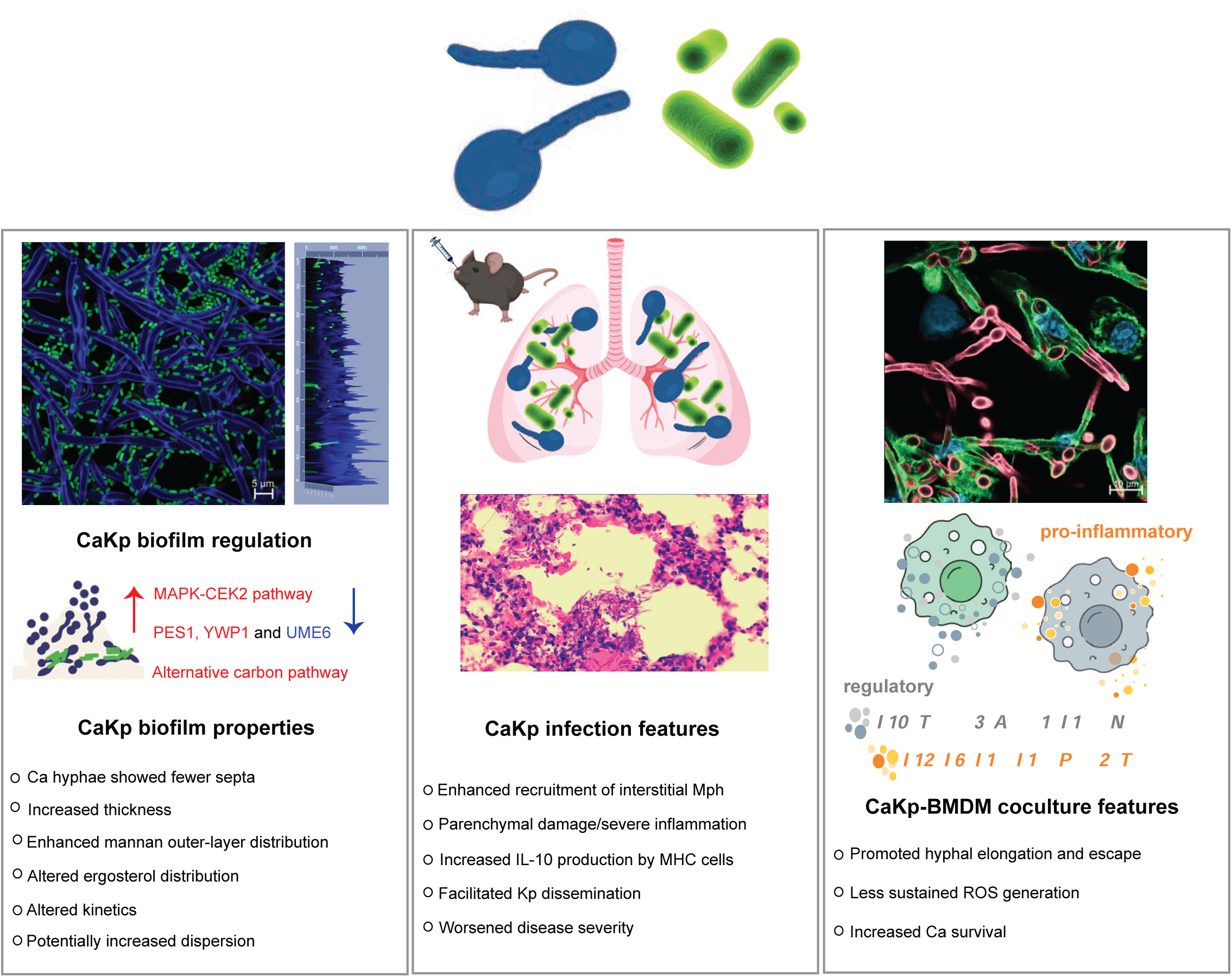
Integrated overview of *Candida–Klebsiella* interactions across *in vitro* and *in vivo* models. Experimental systems include *in vitro* mixed-species biofilms, *in vivo* acute fungal pneumonia, and a macrophage infection model used to investigate host–pathogen interactions. *In vitro* biofilms show differential regulation of biofilm dispersal genes (*PES1, YWP1,* and *UME6*), the cell wall integrity (CWI) MAPK–CEK pathway, and activated the alternative carbon metabolism. Induced and repressed pathways/genes are shown in red and blue, respectively. Mixed-species biofilms exhibit increased biofilm thickness, reduced hyphal septation, and altered cell wall composition, including changes in ergosterol content and mannan distribution. *In vivo* co-infection with *C. albicans* and *K. pneumoniae* is associated with bacterial dissemination, increased macrophage recruitment, and enhanced hyphal invasion into lung tissue, resulting in marked tissue damage. In bone marrow–derived macrophages (BMDMs), co-culture induces both inflammatory (*Il12, Il6, Il1b, Ptgs2,* and *Tnf*) and regulatory (*Il10, Tnfaip3, Acod1, Il1rn,* and *Nfkbia*) responses, shown in orange and grey, respectively, alongside reduced ROS dynamics compared to single infection, with enhanced *C. albicans* survival.

## DISCUSSION

Fungal morphogenesis is a widely accepted determinant of *C. albicans* virulence^6,7,38^. Hyphal elongation and filamentation also facilitate fungal escape from macrophages by breaching phagosomal membranes, ultimately enabling macrophage killing and fungal egression^20^. However, this process is actively sensed by host cells and triggers compensatory immune responses, including modulation of inflammatory programs ^39–41^. It has also been shown that cell wall remodeling programs in phagocytosed *C. albicans* can rapidly attenuate the pro-inflammatory state of host cells, potentially through masking of fungal recognition via alterations in outer cell wall architecture, involving particularly changes in mannan carbohydrate composition ^42,43^.

Bacterial-induced fungal cell wall remodeling has been demonstrated in polymicrobial settings, and recent studies have begun to elucidate how such interactions shape immune recognition and responses ^44,45^. Notably, gastrointestinal bacteria, including *Escherichia coli*, can modulate fungal β-glucan exposure by altering both mannan content and architecture, as well as the spatial organization of β-glucans, thereby influencing phagocytosis efficiency^13^.

Consistent with these findings, our data support a model in which bacterial–fungal interactions drive extensive regulation of cell wall remodeling and mannan redistribution, accompanied by pronounced rewiring of hyphal morphogenesis programs promoting fungal filamentation. These morphogenetic changes are coupled to elevated cell wall stress signaling pathways and shifts in carbon metabolic routes. Strikingly, rapid hyphal elongation in a *Candida–Klebsiella* co-infection setting coincides with a tightly regulated macrophage response characterized by a strong proinflammatory signature alongside an anti-inflammatory response, leading to impaired fungal containment and increased tissue damage.

*C. albicans* and *K. pneumoniae* are clinically important pathogens that represent a significant global public health threat ^24,46^. These organisms can co-occur in the respiratory tract and potentially interact with each other, as reported in cases of COVID-19 patients, where they were among the main etiological agents of infectious complications^47^. Indeed, *Candida* spp. colonize the respiratory tract alongside bacterial communities in a substantial proportion of both healthy individuals and patients with respiratory diseases, including asthma, COPD, cystic fibrosis, and bronchiectasis^48^. The transition from colonization to infection is strongly influenced by microbiome dynamics as well as immune dysregulation. Notably, antibiotic-induced dysbiosis in murine models promotes *C. albicans* expansion, leading to altered Dectin-1 expression in alveolar macrophages, enhanced inflammatory responses and lung damage, exacerbating *K. pneumoniae*-associated infection^49^.

From a clinical diagnostic perspective, the detection of *C. albicans* alongside bacterial respiratory pathogens remains challenging, and therapeutic decisions are often a double-edged sword, as antibiotic use may promote fungal overgrowth and antifungal treatment can mediate selection of resistant strains. Despite these observations, it remains unclear whether such co-occurrence is merely incidental or reflects biologically meaningful interactions enhancing pathogenicity and contributing to impaired disease outcomes. These complexities highlight the need for a deeper mechanistic understanding of polymicrobial interactions. Accordingly, this study was directed to investigating how *K. pneumoniae* reshapes *C. albicans* physiology, virulence-associated traits, and host responses during co-infection, and how these interactions contribute to disease severity.

Here, we show that in response to *K. pneumoniae* encounter, *C. albicans* rewires carbon metabolism, enhances cell wall integrity (CWI) and activates MAPK–Cek2 pathways. Concomitantly, the fungus reduces cell cycle progression and septation, processes that act in concert to favor hyphal elongation^29^. Cell wall remodeling and fungal morphogenesis are highly dynamic and tightly regulated processes that arise from coordinated crosstalk between multiple signaling pathways responding to diverse environmental cues, including nutrient availability^50^. Consistent with this, our data indicate that the response of *C. albicans* to *K. pneumoniae* differs between M9 and RPMI media, likely reflecting differences in metabolic state and growth dynamics.

Biofilm formation by *C. albicans* is governed by a core transcriptional network of genes comprising *BCR1, TEC1, EFG1, NDT80, ROB1,* and *BRG1*^51^. In our dataset, CaKp biofilms showed broad activation of this network under M9 conditions, with upregulation of at least five genes, whereas in RPMI only *ROB1* showed strong induction. This differential activation likely reflects nutrient-dependent biofilm dynamics. Under nutrient-limited M9 conditions, where growth is constrained, transcriptional programs associated with initiation of hyphal development and early biofilm structural organization appear to be preferentially engaged, suggesting delayed biofilm formation kinetics. In contrast, in RPMI, a condition more permissive for growth and filamentation, regulatory activation may instead reflect progression toward later stages of biofilm development, characterized by distinct transcriptional programs involving factors such as *UME6* and *YWP1*.

Considering the biofilm development in RPMI, our data further revealed the impact of fungal-bacterial interaction on *Candida* biofilm thickness already at early time points. Moreover, the mixed biofilms showed changes in septation and redistribution of outer-layer mannan as well as marked upregulation of *YWP1,* as highlighted above. *YWP1* encodes a highly abundant cell wall mannoprotein implicated in yeast cell dispersal from biofilms and in β-1,3-glucan masking^52^. Alterations in cell wall composition such as in β-glucan exposure are strongly correlated with changes in cytokine responses and phagocytosis efficiency^53,54^. Our data further corroborate these observations by showing that CaKp co-infection modulates the host immune response, as reflected by cytokine secretion signatures by BMDMs, with increased release of TNF-α, IL-10, and MCP-1. Moreover, the interaction affected the extracellular fungal burden, likely reflecting reduced phagocytic uptake and enhanced fungal survival. Concurrently, co-infection impairs sustained macrophage ROS production and oxidative burst activity, thereby facilitating fungal immune evasion.

*In vivo*, co-infection exacerbated lung tissue damage, coinciding with extensive *C. albicans* filamentation and slightly increased bacterial dissemination, reflecting the enhanced pathogenicity of the microbial co-infection. In line with this finding, *K. pneumoniae* is known for being able to create an anti-inflammatory metabolic milieu in lung tissues that promotes bacterial persistence^55^. In the context of our model, this may also generate conditions permissive for hyphal elongation and fungal escape. Additionally, *ex vivo* restimulation of lung leukocytes with heat-inactivated pathogens revealed enhanced IL-10 production in co-infected mice, a response largely absent in the *K. pneumoniae*-primed group. Together, these data support the concept that, during co-infection, macrophages adopt a continuum of activation states characterized by overlapping transcriptional programs. This response includes an initial pro-inflammatory phase, evidenced by induction of *Ptgs2* and *Tnf*, followed by engagement of a type I interferon–associated regulatory module, including *Il10*, *Il1rn*, and interferon-stimulated genes such as *Rsad2* and *Ifit2*. This coordinated response suggests that type I IFN signaling does not merely amplify inflammation but rather integrates feedback mechanisms that help limit excessive tissue damage. The resulting mixed activation state may therefore create a partially permissive environment that favors *C. albicans* persistence.

Previous studies have shown that *C. albicans* co-culture with macrophages exposed to bacterial components, such as lipopolysaccharide, can regulate immune responses with enhanced secretion of IL-10, in a morphotype fungal-dependent manner^56^. We further propose that bacterial metabolism and cross-feeding within the macrophage environment may actively shape the immune landscape, generating conditions permissive for polymicrobial persistence and enhancing *C. albicans* immune evasion. In line with this, our data indicate that tryptophan metabolism is altered in bacterial cells during co-infection. Trypthophan metabolism and indole-derived metabolites produced by *K. pneumoniae* function as ligands for activation of the aryl hydrocarbon receptor (AhR) pathway in the host. AhR signaling has been linked to immunoregulatory programs, including IL-10 production^57^. Herein, we further demonstrate engagement of this pathway in response to *Klebsiella* with marked upregulation of *Tiparp*, *Ido2* and *Ahr*.

Taken together, our data indicate that *C. albicans–K. pneumoniae* co-infections prompt the host immune defense to establish a tightly regulated innate phagocyte environment in response to both microbial pathogens. Within this scenario, *Candida* exploits the permissive niche by enhancing escape and hyphal development, while *K. pneumoniae* benefits from the well-developed hyphae to promote dissemination Understanding the regulatory mechanisms that govern this *Candida*-driven adaptation will be critical for developing strategies to mitigate the severity of mixed bacterial-fungal infections.

## Limitations of the study

Possible limitations arise when our *in vitro* transcriptomic and bone marrow–derived macrophage (BMDM) findings are extrapolated to the pulmonary infection context *in vivo*. The lungs are a highly specialized organ with restricted nutrient-accessibility and a complex cellular immune milieu comprising resident and recruited immune as well as non-immune cells, including alveolar and interstitial macrophages, epithelial cells, dendritic cells, innate lymphoid cells, neutrophils, and diverse lymphocyte populations. Our biofilm RNA-seq analyses were performed in RPMI and M9 media, supporting growth and biofilm formation mainly under RPMI conditions for both species. However, the culture conditions did not fully mimic the metabolic constraints or cellular complexity within the pulmonary environment. Similarly, our RNA-seq analyses focused on primary BMDMs, which share developmental and functional features with recruited interstitial macrophages but do not capture the transcriptional and metabolic specialization of alveolar macrophages. Thus, these systems represent valid but reductionist models to address mixed biofilm interactions and cell-intrinsic macrophage effector functions in antimicrobial defense. Despite these limitations, macrophages comprise a large population of immune cells affecting lung homeostasis and supporting the relevance of a macrophage-centered approach. Moreover, *in vivo* measurements of fungal burden, immune cell composition by flow cytometry, and histopathology strongly support the biological relevance of our findings. Nonetheless, conclusions regarding metabolic crosstalk and niche-specific responses should be interpreted within the context of these experimental constraints.

## RESOURCE AVAILABILITY

### Lead contacts

Further information and requests for resources and reagents should be directed to and will be fulfilled by the lead contacts, Karl Kuchler and Thomas Lion.

### Materials availability

Further information and reasonable requests for resources and reagents should be directed to and will be fulfilled by lead contacts.

### Data and code availability

RNA-seq data reported in this manuscript have been deposited in the GEO databases (GSE292174, GSM8811286). Data reported are available from the lead contact upon request. Any additional information required to re-analyse the data reported in this paper is available from the lead contact upon request.

## ACKNOWLEDGMENTS

We thank Andriy Petryshyn and all lab members for technical support and experimental advice, and the Max Perutz Labs Vienna (MPL) Mouse Facility for maintaining the mouse lines. We also acknowledge the service of the MPL Histology Facility and MPL Optics Facility. We also would like to thank Emir Hadzijusufovic and Peter Valent for scientific advice. We also acknowledge Leonel Pereira and Shirin Sharghi for their early involvement in the project. This work was supported by grants from the Austrian Science Fund FWF (FWF-SFB and BacFun to T.L. and K.K.).

## AUTHOR CONTRIBUTIONS

Conceptualization, T.B., I. T., K.K, and T.L.; methodology, T.B. I.T. and R.F.S.; investigation, T.B., I.T., R.F.S, N.W., T.P.-C., C.P, S.V.P., S.H TZ, PP; writing—original draft, T.B., I.T.; writing, review & editing, K.K., T. L., W.E., M. L., D. C-H., T.B., I.T., R.F.S.; visualization, T.B., I.T., T.P.-C; funding acquisition, K.K., TL.

---

**Extended Figure 1.**
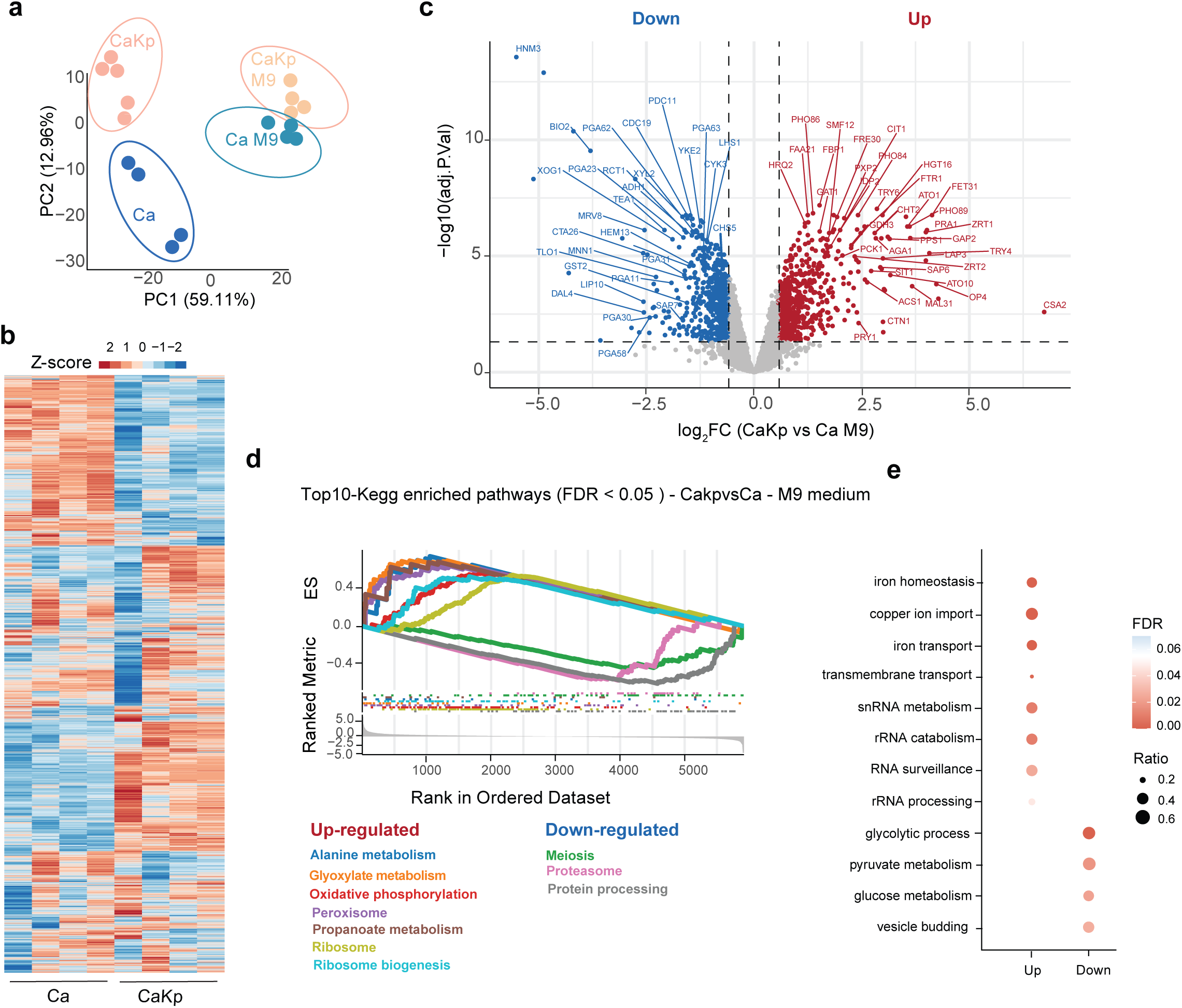
Transcriptional dynamics of *C. albicans* in mixed-species biofilms with *K. pneumoniae* grown in M9 medium. **(a)** Principal component analysis (PCA) of transcriptomes from single-species *C. albicans* cultures (Ca) and co-cultures with *K. pneumoniae* (CaKp) grown in M9 and RPMI media, showing a clearer separation between conditions in RPMI. **(b)** Hierarchical clustering (Euclidean distance) of gene expression across all genes following Trimmed Mean of M-values (TMM) normalization. Expression values are shown as log counts per million (logCPM) and scaled as Z-scores. Four biological replicates are shown for Ca and CaKp cultures grown in M9. **(c)** Volcano plot of differentially expressed genes (DEGs) comparing CaKp versus Ca in M9, plotted as log2 fold change (log2FC) versus −log10 adjusted *P* value. DEGs were defined using a log2FC cutoff of ±0.5. Selected genes are indicated. **(d)** Gene set enrichment analysis (GSEA) of KEGG pathways for *C. albicans*. Pathways with Benjamini–Hochberg false discovery rate (FDR) < 0.05 were considered significantly enriched. **(e)** Gene Ontology (GO) enrichment analysis of biological processes associated with upregulated and downregulated DEGs in CaKp compared to Ca.

**Supplementary Figure S1:**
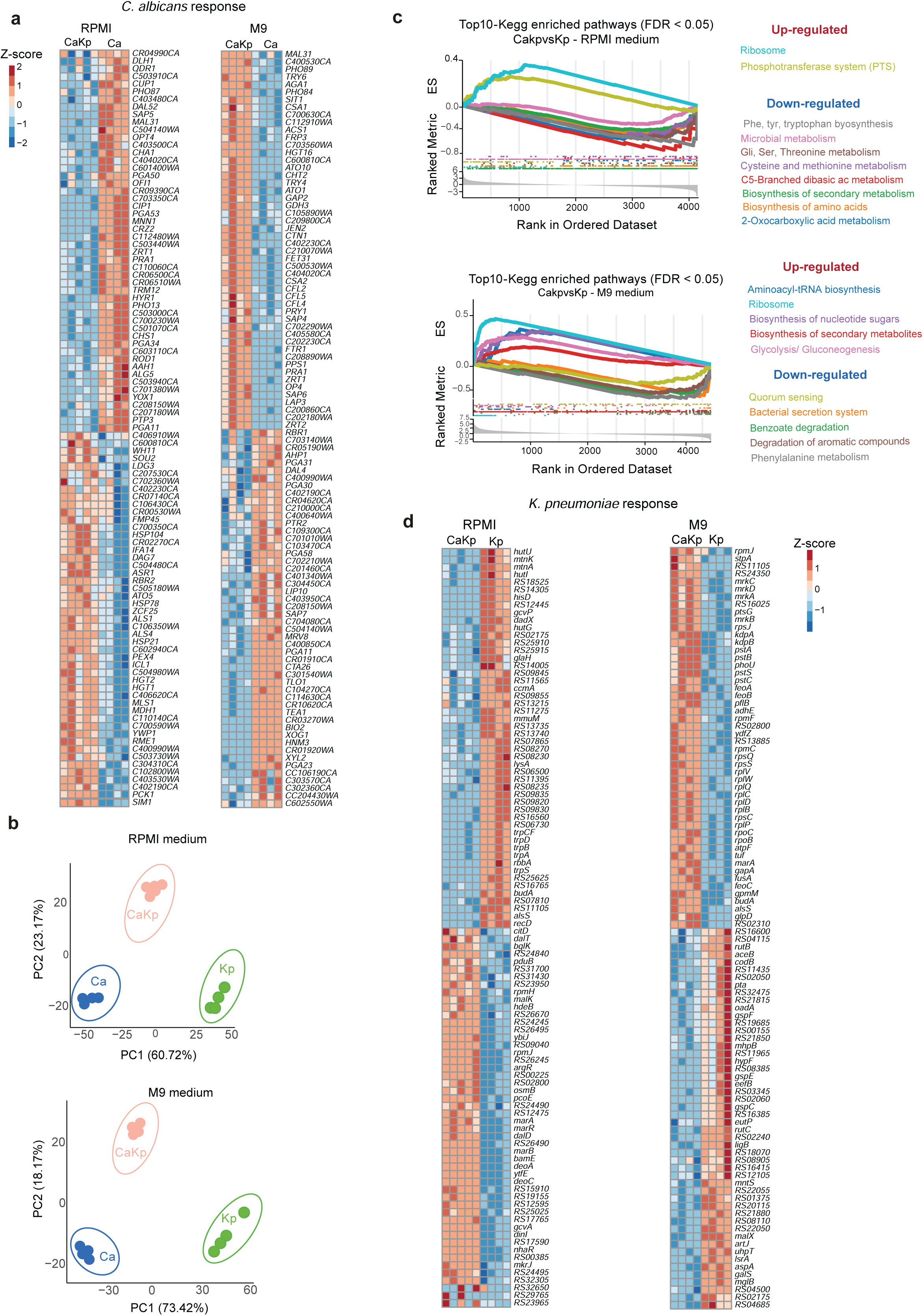
Transcriptional responses of *C. albicans* and *K. pneumoniae* in mixed biofilms. **(a)** Top expressed genes in single and co-culture conditions, illustrating transcriptional responses of *C. albicans* in RPMI and M9 media. **(b)** Principal component analysis (PCA) of transcriptome profiles from Ca, Kp and CaKp grown in RPMI (top) and M9 (bottom). **(c)** Gene Set Enrichment Analysis (GSEA) of KEGG pathways in *K. pneumoniae* from biofilms grown in RPMI (top) and M9 (bottom). Pathways with *P* < 0.05 were considered significantly enriched**. (d)**Top expressed genes in single and co-culture conditions, illustrating transcriptional responses of and *K. pneumoniae* in RPMI and M9 media.

**Extended Figure 2.**
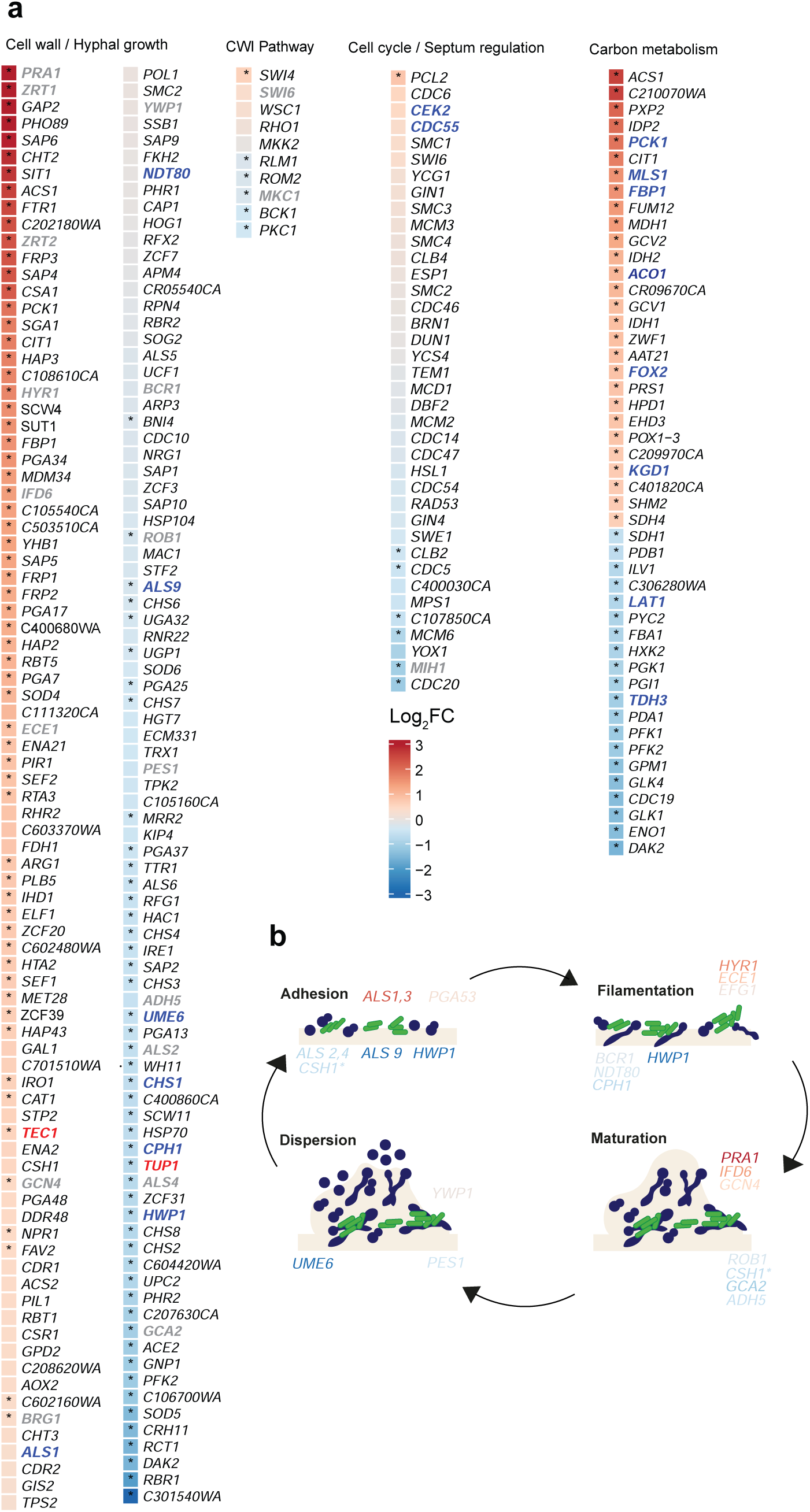
Pathways regulated during the *C. albicans–K. pneumoniae* interaction and impact on *Candida* biofilm formation in M9. **(a)** DEGs in selected categories, including hyphal growth/cell wall, CWI pathway, cell cycle, and carbon metabolism, comparing CaKp *versus* Ca. Color scales represent log2-fold change (log2FC). Genes highlighted in blue are consistently regulated in *C. albicans* regardless of growth medium (RPMI or M9); genes in grey indicate medium-dependent responses with differential behaviour between RPMI and M9; and genes in red denote those involved in hyphae and biofilm formation exclusively modulated in M9. **(b)** Schematic representation of biofilm maturation stages, and DEGs potentially involved in each stage after *Candida-Klebsiella* interaction.

**Figure S2.**
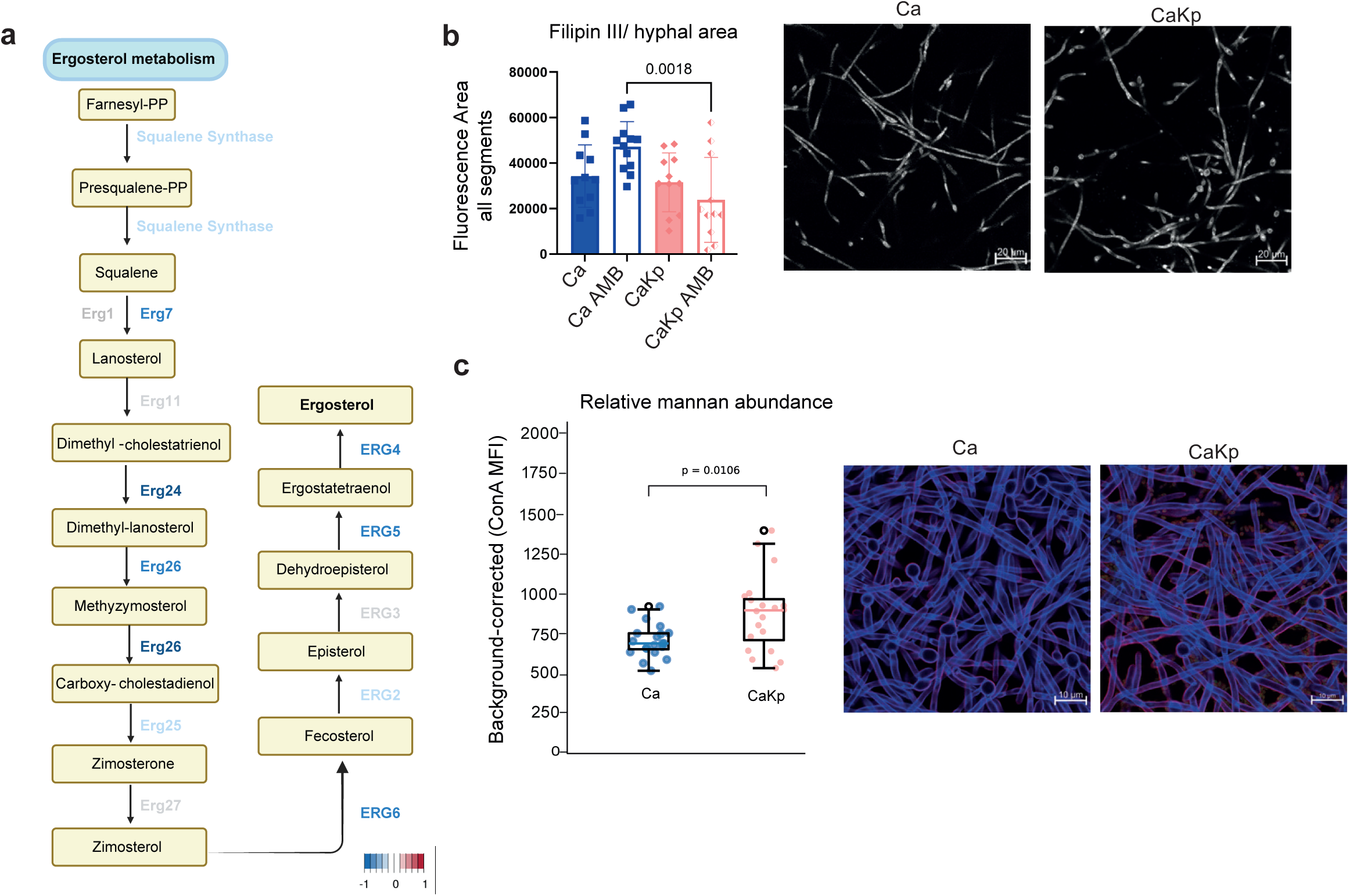
*Klebsiella* reduces expression of ergosterol biosynthesis genes in *C. albicans*, as reflected by altered filipin distribution, and increases in mannan content. (a) Ergosterol biosynthesis pathway showing gene expression modulation at each step (adapted from KEGG). **(b)** FilipinIII staining of *C. albicans* biofilms at 24h time point, and quantification of fluorescence intensity comparing *C. albicans* (Ca), Ca exposed to Amphotericin B (AMB) 0.5 μg/ml for 4.5h, *C. albicans* with *K. pneumoniae* (CaKp) and CaKp exposed to AMB 0.5 μg/mL for 4.5h. **(c)** Total Mean of fluorescence of Concanavalin A (ConA) staining indicating relative mannan content.

**Figure S3.**
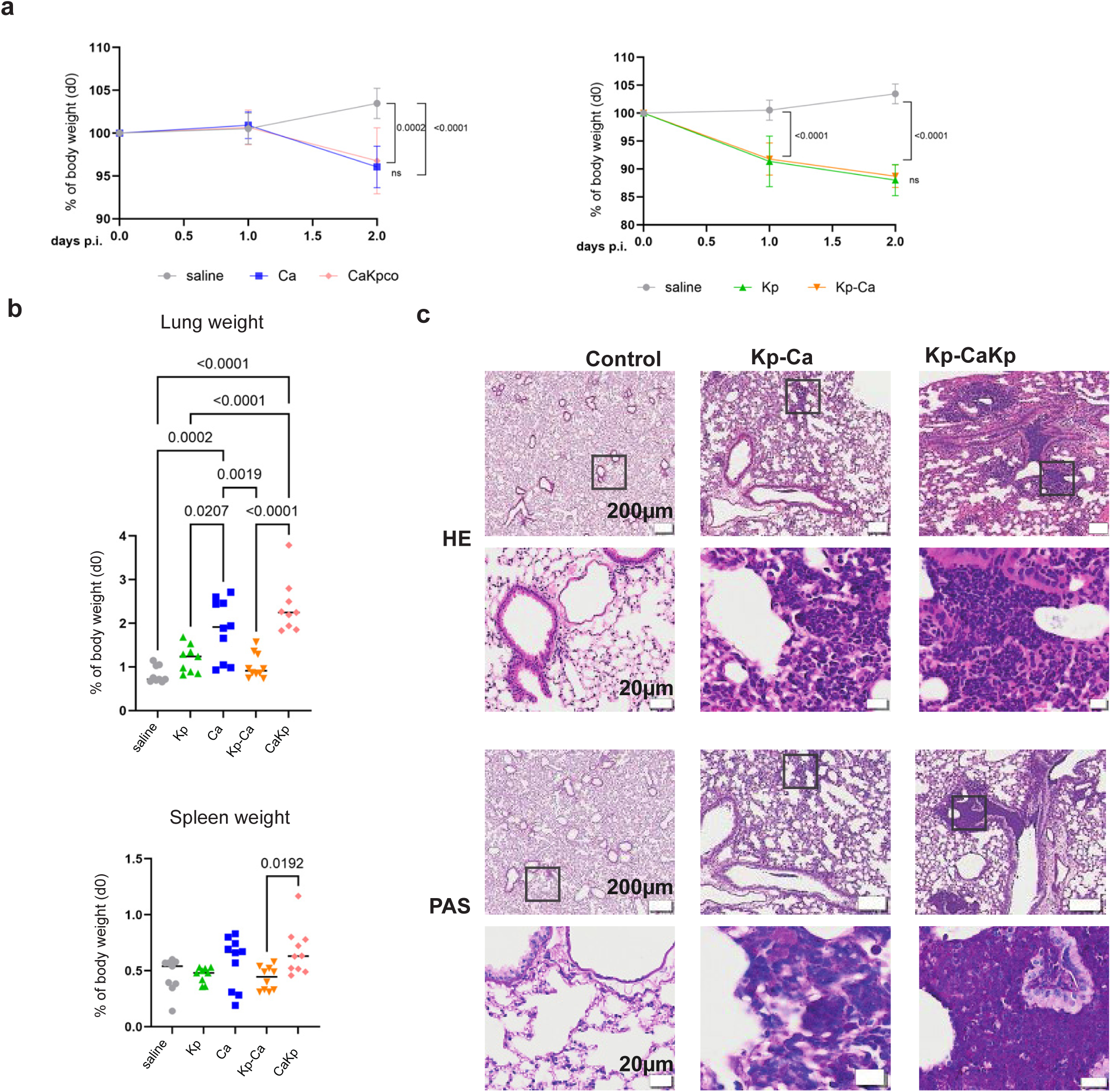
Impact of single and co-infection in mice on organ weights and lung pathology, and effect of *Klebsiella pneumoniae* priming. **(a)** Changes in mouse body weight following mono- and co-infection (n ≥ 9 animals from three independent experiments). Left, saline controls and *Ca*-infected mice; right, saline controls and *Kp*-infected mice.. **(b)** Lung and spleen weights. **(c)** HE and PAS of lungs primed by Kp challenge, followed by the infection with *Candida* or *Candida-Klebsiella*. Scale bars, 20 μm and 200 μm.

**Figure S4.**
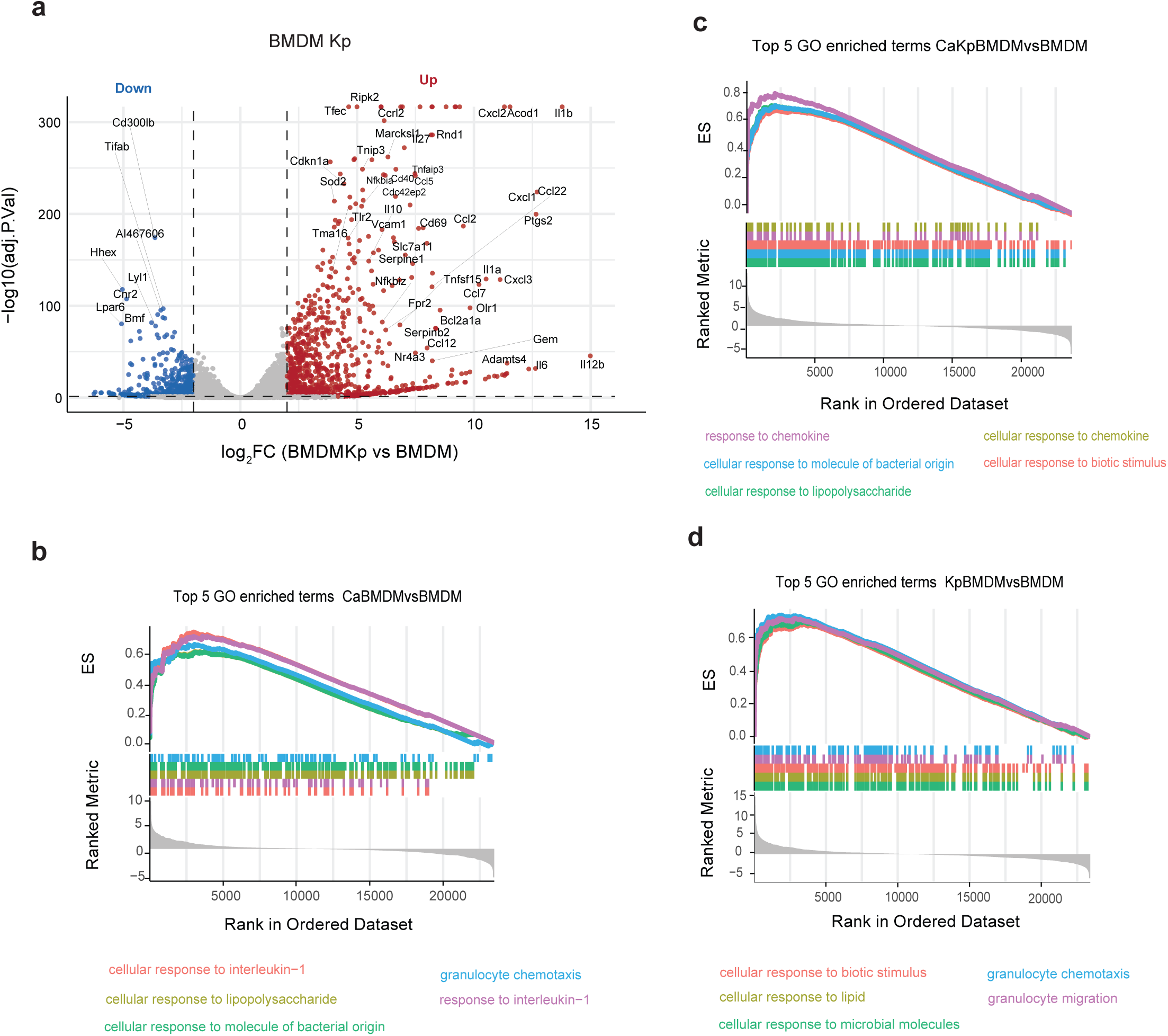
Transcriptional response of BMDMs to *K. pneumonia (Kp)*, *C. albicans (Ca)* and CaKp). **(a)** Volcano plot representing the up- and down-regulated DEGs from the comparison KpBMDM *vs* BMDM, using log2-fold change (log2FC) *versu*s −log10-adjusted P value for all genes. **(b)** GSEA covering GO biological processes terms from CaBMDM *vs* BMDM or CaKpBMDM *vs* BMDM **(c)**. Pathways with adjusted P value (Benjamini–Hochberg FDR) < 0.05 were as considered significantly enriched. Only upregulated processes are shown. **(d)** GSEA covering the GO biological processes terms form the comparison KpBMDM *vs* BMDM. Pathways with adjusted P value (Benjamini–Hochberg FDR) < 0.05 were considered as significantly enriched. Only upregulated processes are shown.

**Figure S5.**
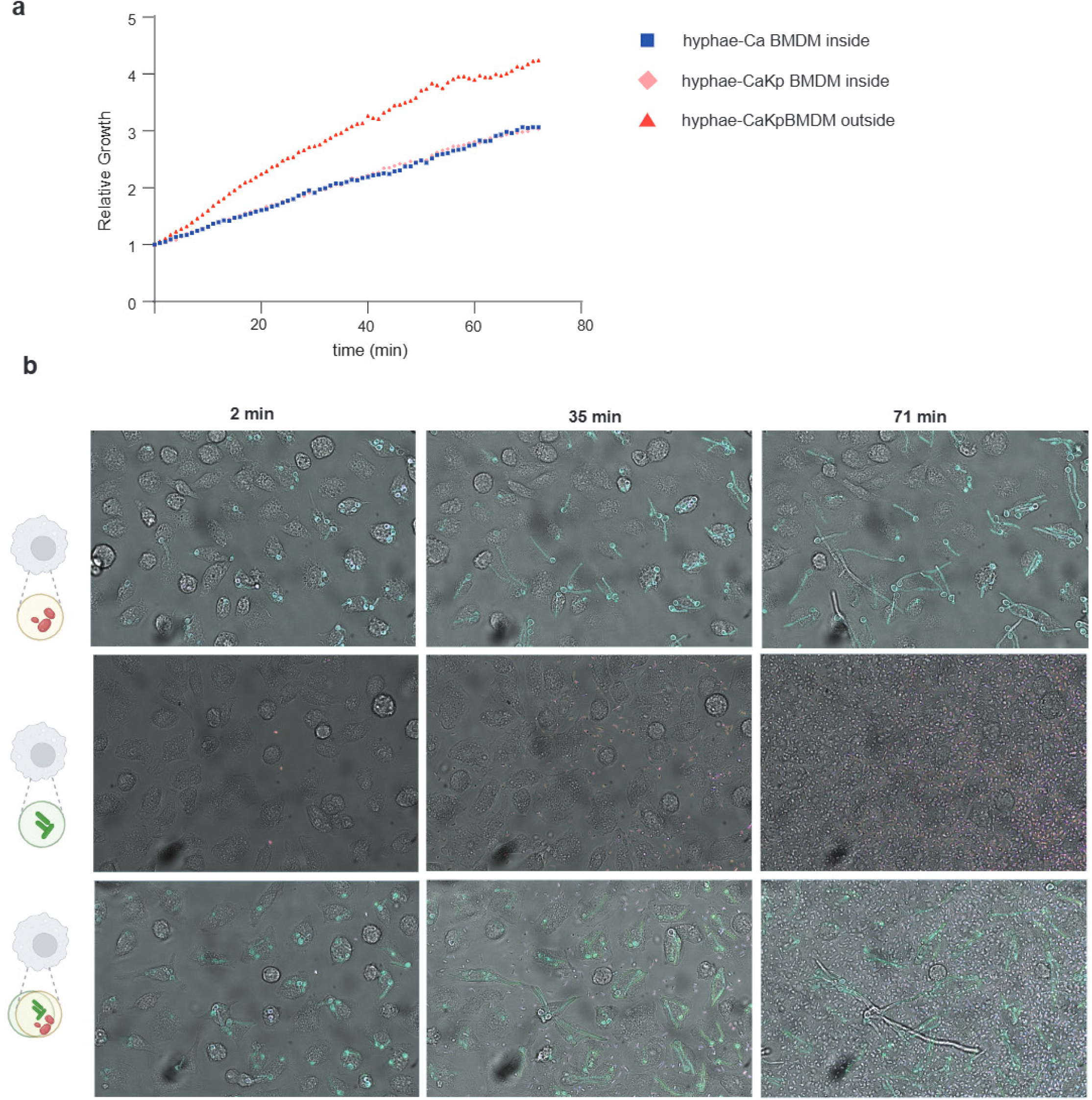
Co-culture with *Klebsiella* enhances hyphal growth and macrophage escape by *C. albicans,* as shown by live-cell imaging over 80 min. (a) Growth curve showing increased hyphal elongation and escape from BMDMs under CaKp co-culture condition compared with Ca alone. **(b)** Representative time-lapse images of BMDMs exposed to Ca-dTom, Kp-GFP, or CaKp, captured by live-cell imaging at different time points.

## EXPERIMENTAL MODEL DETAILS

### Ethics statement and animal housing

All animal experiments were performed in accordance with institutional regulations and approved by the ethics committee of the Medical University of Vienna and the Austrian Federal Ministry of Education, Science and Research (BMWFW-66.009/0436-V/3b/2019). Mice were housed under specific and opportunistic pathogen-free (SOPF) conditions in individually ventilated cages (IVCs) at 20-25 °C, 45-65% relative humidity and 14 h light/10 h dark cycle. Food and water were provided *ad libitum*. C57BL/6J female mice were bred and maintained at the Max Perutz Labs Vienna.

### Strains and growth conditions

*Candida albicans* SC5413 and *K. pneumoniae* ATCC 700603 are the strains used in this study. *C. albicans* was maintained in yeast extract peptone dextrose (YPD) medium (10 g/L yeast extract, 20 g/L peptone, 2% glucose) (Formedium, Norfolk, UK), growing for 2 days at 30 °C. Then *C. albicans was* pre-cultured at 30°C with agitation at 200 rpm; cells were washed two times with phosphate buffered saline (PBS). *K. pneumonia* strains were maintained in Luria-Bertani (LB) agar plates, and precultured for 12–16 h in liquid LB medium at 37 °C with agitation at 180 rpm. An equal cell numbers of *C. albicans* and *K. pneumonia* (10^6^ cells/mL) were used in co-culture experiments. For transcriptomic and confocal microscopy, single pathogen cultures and co-cultures of *C. albicans* and *K. pneumoniae* were grown as biofilms, with static incubation either in treated 6-well plates (CytoOne, Starlab # CC7682-7506) or 35 × 10 mm biTreat culture dishes (Ibidi #81156), respectively. Growth was allowed for 1.5h for adherence followed by 24h at 37 °C in 3 mL medium RPMI (Corning, # Cat#COR50-020-PC) supplemented with 2% glucose for RNA-seq, or 1.5 h for adherence followed by 6h in case of confocal images. To assess the effect of nutrient availability on biofilm dynamics, an additional set of single- and mixed-species biofilms was included in the transcriptomic experiments. These biofilms were grown in M9 medium (M9 salts, Sigma) supplemented with 0.4% glucose, 0.002 M MgSO₄, and 0.0001 M CaCl₂.

### Isolation and differentiation of bone marrow-derived macrophages (BMDMs)

BMDMs were isolated and differentiated from bone marrow precursors, as previously described. ^58^ Briefly, femur and tibia from 8-15-week-female mice (WT C57BL/6J) were collected and cut open on one epiphysis. The pluripotent cells were recovered by centrifugation of the cut bones placed into a 0.5 mL tube inserted in 1.5mL tube for 2min at 13,000 rpm. Thereafter the cells were filtered in a 40µm strainer (Corning # 352340) and seeded in macrophage medium, containing Dulbecco’s modified Eagle’s medium – DMEM (Gibco # 11995065), 10% heat-inactivated fetal bovine serum – hiFBS (Sigma #F7524), 15% L929-supplement, and 1 x Penicilin/Streptomicin (#15140-122). During the differentiation and propagation of BMDMs, half of the media was renewed at day 3, and the cells were split at day 5, and at day 7, the BMDMs were harvested and seeded accordingly to assay purposes.

## METHOD DETAILS

### Mouse intranasal infection model

*Klebsiella pneumoniae* and *Candida albicans* were pre-cultured at 30°C and 37°C, respectively, with agitation at 200 rpm until the exponential growth phase and diluted in sterile, physiological saline (Fresenius) to final concentrations of 1x10⁷ CFU and 5x10⁶ CFU per 40µL, respectively. For co-infection experiments, pathogens were mixed at a 2:1 ratio, resulting in 5x10⁶ CFU *K. pneumoniae* and 2.5x10⁶ CFU *C. albicans* in 40 µL. Mice were anesthetized via intraperitoneal injection of ketamine and xylazine. 40 µL of the pathogen suspension was administered intranasally and taken up by spontaneous inhalation. Mock-infected control animals received 40 µL of sterile saline. Infections or saline treatments were repeated the following day as indicated. Mice were monitored daily for weight loss and clinical appearance. Humane endpoints were established in accordance with institutional animal care guidelines, federal regulations and FELASA recommendations. To determine dissemination and pathogen burden within tissues, lungs and spleens were aseptically excised, placed in 1 mL sterile PBS (#D8537, Sigma-Aldrich) and weighed. Spleens were mechanically homogenized using an IKA T10 basic Ultra-Turrax homogenizer (IKA, Staufen) under sterile conditions. Lung tissue was enzymatically digested prior to homogenate collection. Serial dilutions of tissue homogenates were plated in triplicate on YPD agar supplemented with ampicillin (120 µg/mL), tetracycline (20 µg/mL), and chloramphenicol (40 µg/mL). Colonies were enumerated, and fungal burden was calculated as CFU of *C. albicans* per gram of tissue.

### Isolation of lung immune cell populations

All procedures were performed under sterile conditions in a laminar flow hood (Safe Fast Premium 212). Mice were euthanized by cervical dislocation, and lungs were immediately harvested into 15ml tubes (Aptaca, #10450/SG) containing 1 ml DPBS (Sigma-Aldrich, #D8537) and placed on ice. Lung tissue was minced into small fragments and digested in Hank’s Balanced Salt Solution (HBSS; Sigma-Aldrich, #H8264) supplemented with 2% heat-inactivated fetal bovine serum (hiFBS; Sigma-Aldrich, #F7524), 0.2 mg/mL DNase I (Roche, #10104159001), and 1 mg/mL Collagenase II (Gibco, #17101015). Digestion was carried out for 30 minutes at 37 °C with agitation (109 rpm, 40° angle) in 50 ml tubes (Starlab, #E1450-0200) and terminated by transferring samples to room temperature (22–24 °C). Cell suspensions were filtered through a 70 µm cell strainer (Falcon) into ice-cold FACS buffer (HBSS containing 2% hiFBS and 2.5 mM EDTA). An aliquot of the suspension was used for fungal burden quantification. Red blood cells were lysed using Red Blood Cell Lysis Buffer (BioLegend, #420302), followed by density gradient centrifugation (40%/80% Percoll; GE Healthcare, #GE17-0891-01) at 1000xg for 20 min at 22 °C in 15 mL tubes (Starlab, #E1415-0200). The interphase was collected and washed in FACS buffer. Total cell counts were determined using a CASY automated cell counter (OLS) with a size threshold set between 5–20 µm. Cells were aliquoted for downstream applications: 1.5x10⁶ cells for flow cytometry staining and 3x10⁶ cells for *ex vivo* restimulation. Restimulation was performed in 96-well plates with DMEM supplemented with 10% hiFBS, ampicillin (100 µg/mL; Sigma-Aldrich). Cells were incubated together with heat-inactivated pathogens at a MOI of 1 for 5.5 h at 37°C and 5%CO_2_. Brefeldin A (BioLegend) was added 2 h after stimulation onset to promote intracellular cytokine accumulation.

### Histology

Mice were euthanized on day 3 post-infection by intraperitoneal overdose of ketamine (20 mg; Livisto) and xylazine (0.4 mg; Dechra), followed by transcardial perfusion with 15 mL PBS and two subsequent washes with 15 mL of 4% paraformaldehyde (PFA; #11773266, Liquid Productions) in PBS. Lungs were inflated with 4% PFA via intratracheal instillation and subsequently incubated at 4°C in 5 mL of 4% PFA for fixation. After 24 h, the fixative was replaced with PBS, and tissues were processed for paraffin embedding. Paraffin-embedded lungs were sectioned at 4 µm thickness using a microtome (Leica) and stained with hematoxylin and eosin (H&E) or periodic acid–Schiff (PAS) using standard histological protocols. Blinded scoring of inflamed areas on stained sections was performed by a certified pathologist. Representative images were acquired using an Olympus BX61VS microscope equipped with an automated slide scanning stage and OlyVIA software (version 2.9; Olympus).

### *In vitro* survival assay

For survival assay, procedures were carried out as previously described^59,60^ with few changes. Briefly, 1x 10^5^ BMDMs were seeded into tissue culture-treated 96-well plates (Starlab), in 100µl of macrophage medium without antibiotics, and kept for 24h for adherence, then *C. albicans* and *K. pneumoniae* were added at the multiplicity of infection (MOI) of 0.1. The co-interaction of immune cells with the pathogens were kept for 3 h at 37 ^0^C with 5% CO_2_. After 3h incubation time, the plate was put on ice, and then added 50ul of a 4% Triton-X 100/PBS solution, pipet up and down and cells were then recovered by scratching the wells with a tip, followed by 2 times washing with PBS. The cells were collected in 400ul PBS tube. Then the collected cells were plated in extract-peptone-glucose (YPD) agar and kept at 30 degrees for 48 h for CFU counts.

### Phagocytosis assays

The phagocytosis assay was performed as previously described^61^ with few changes. Briefly, 5x10^5^ BMDMs were seeded in 24-well plate (Starlab) in 500 µl macrophage medium without antibiotics (DMEM with 10% FBS, and 15% of L929), and incubate at 37°C with 5% CO_2_ for 24h. Thereafter, the cells were co-infected with 50µl labelled *C. albincans* dTom, *K. pneumoniae* GFP and with both pathogens concomitantly at a MOI: 2 (1 immune cell: 2 yeast cells: 2 bacterial cells). Plates were then incubated at 37 °C with 5% CO_2_ for 45min and 120min. After the incubation time, the supernatant (SN) was recovered for further analysis of non-phagocyted *Candida/Klebsiella*. Then, the cells were washed 3 times with PBS, and recovered after treatment with 250 μl trypsin (Sigma-Aldrich). BMDMs were harvested after adding 750 μl DMEM-10% hiFBS and then detaching cells by pipetting followed by centrifugation at 400×g at 4°C for 4 minutes. Cells were then resuspended in 50 μl of FACS buffer (PBS w/o Ca^2+^ and Mg^2+^ with 2-5% hiFBS, 5mM EDTA), and then resuspended in FACs buffer containing CD16/32 (1:1000, Biolegend #101302), then kept at 4 °C for 5min, followed by the incubation with a rabbit anti-Candida serum^62^ (1:500), and F4/80(1:100, Biolegend, #123116), and then incubated for 15 min at 4°C. The tubes were then filled up with FACs buffer and centrifuge at 400g for 4min at 4°C. Then the cells were recovered and incubated with the secondary anti-rabbit antibody TexaRedaRabbit. (1:200, Invitrogen #T2767) 0,33ul/50ul, and the the viability antibody violet (1:500, Biolegend # B423108), followed by incubation for 15min at 4°C. Then the tubes were filled up with FACs buffer and followed by centrifugation at 400×g at 4°C for 4 min, and the cells were then resuspended in 300ul FACs buffer. The phagocytosis were then evaluated with an LSRFortessa cytometer (BD Biosciences). Data were analyzed with FlowJo v10.8 software.

### Reactive oxygen species (ROS) assays

The ROS assay was carried out as previously described^63^ BMDMs were seed 5x 10^4^ per well into a white 96-well luminometer plate (Thermo Scientific Nunc # 136101) in macrophage medium without antibiotics, and kept 12-16h for adherence. Then, the medium was removed and the cells were washed twice with Hank’s balanced salt solution (HBSS; with Mg2+ and Ca2+; GIBCO #14025-050), and then finally mixed with 50 uL HBSS containing 200 mM luminol (Sigma # A4685) and 16 U horseradish peroxidase – HRP type IV (Sigma, #P8375-5KU). Immediately afterward, pre-cultured C*. albicans and K. pneumonia* was added in 50ul of HBSS at a MOI of 5. As controls, As controls, BMDMs without stimulation, *C. albicans* alone, *K. pneumonia* alone, heat-killed *C. albicans* and *K. pneumoniae*, and LPS were included. Chemiluminescence was monitored on a Victor Nivo microplate reader (PerkinElmer) at 37°C. The detected relative luciferase units (RLU) are expressed as RLU per min per 1000 immune cells or as total RLU after the indicated time.

### Immunofluorescence assays

Immunofluorescence staining was performed as follows. BMDMs were seeded at 10⁵ cells per well (Ibidi, # 80826) and allowed to adhere for 24 h at 37°C with 5% CO₂. Cells were then infected at an MOI of 2 (2 fungal cells and/or 2 bacterial cells per macrophage) and incubated for 2 h at 37°C with 5% CO₂. After infection, supernatants were removed and cells were washed twice with PBS, followed by fixation with 4% paraformaldehyde - PFA (Thermo Scientific, #28906) for 15 min at room temperature (RT). Fixed cells were washed twice with PBS and permeabilized with 0.1% Triton X-100 in PBS (Sigma, #X100) for 15 min at RT. Cells were then washed and blocked with 3% BSA in PBS (Sigma) before incubation with anti-Candida serum (1:500) for 1 h at RT. After two washes with 3% BSA in PBS, cells were incubated with TexaRed anti-rabbit IgG (1:500, #T2767) together with phalloidin (1:50, Cell Signaling Technology, #8878S) for 1 h at RT. Following two PBS washes, DAPI (1:200, # GTX16206-10) was added for 1 min, and cells were washed twice and stored in PBS protected from light.Images were acquired using a Zeiss LSM 980 inverted confocal microscope equipped with an Airyscan detector at the Biooptics Facility of the MPL.

### Live-imaging of cell co-cultures

BMDMs were seeded onto a chambered coverslip (IBIDI, #80826) at a density of 10^5^ cells per well in DMEM without phenol red (Gibo #31053028) supplemented with 25ng/ml murine CSF-1 (Peprotech # AF-CSF-315-02-10UG). dTOM-expressing *C. albican*, GFP-expressing *K. pneumoniae*, or both dTom-*C. albicans* and GFP-*K. pneumoniae* were added at a MOI of 2. Non-infected macrophages were used as a control. Live-cell imaging was performed immediately at 37°C using a Yokogawa W1 Spinning Disk confocal microscope *on a Nikon Ti2-E stand* equipped with a high-resolution **s**CMOS camera (pixel-size 6.5 µm) at the Biooptics facility of the Max Perutz Labs Vienna. Cells were imaged at 40× magnification using a 0.95 NA PlanApo Objective. Fluorophores were excited using a 488 nm laser (dGFP channel) for GFP-*K. pneumoniae*, and a 561 nm laser (dTOM channel) for dTom-*C. albicans,* respectively. Images were acquired with the following parameters: one image every 2 minutes, 200 ms exposure time for each channel. Image acquisition was carried out for a total duration of 4 hours.

Image analysis was performed using a custom Python-based pipeline. Acquired z-stacks were maximum-intensity projected and preprocessed by applying a subtle gaussian filter for noise reduction (σ=2.0), with subsequent gaussian background subtraction. Intensities were normalized per file using the 1^st^ and 99^th^ percentiles. The preprocessed data were then segmented using Otsu’s method. Segmented binary masks were then further refined by morphological closing for bridging small gaps and improving mask continuity, followed by removing small objects below 200 pixels. Masks were then skeletonized using a medial-axis-based skeletonization. Resulting skeletons were pruned to remove short terminal branches shorter than 10 pixels, eliminating spurious branches caused by segmentation noise. Segmentation and skeletonization quality were visually inspected to confirm accurate hyphal detection. Overall growth rates were then quantified by measuring relative increase in overall skeleton length in the entire field of view over time.

### Inflammatory cytokine Elisa panels

Quantification of inflammatory cytokine presence in cell culture supernatant was measured using LEGENDplex Mouse Inflammation Panel kit (Biolegend, #740446). Supernatant was gathered from the BMDMs infected with *C. albicans*, *K. pneumoniae* or both at a MOI of 3 for 2h. Supernatant was recovered and stored at – 80°C until use. Before use, the SN was thawed and centrifuged at 400×g for 5 min to remove cellular debris and then the assay was run as directed by manufacturer’s protocol.

### Total RNA isolation and RNA sequencing from biofilms and BMDMs

Cells from 24 h biofilms of single-pathogen cultures and co-cultures of *C. albicans* and *K. pneumoniae* were collected by centrifugation at 3,000 × g for 15 min at 4°C. The supernatant was removed, and cells were first digested with 0.5 mg/mL lysozyme (Sigma) in 10 mM Tris-Cl, pH 8.0 for 20 min at 37°C, followed by centrifugation at 3,000 × g for 15 min at 4°C. Cells were then resuspended in 1 mL TRI-Reagent LS (LabConsulting) in tubes containing 300 mg glass beads (Sigma-Aldrich) and disrupted by two rounds of bead-beating (FastPrep-MP) at 6 m/s for 45 s, followed by centrifugation at 14,000 × g for 10 min at 4°C. The upper phase was transferred to a fresh tube containing 200 μL chloroform, centrifuged, and the aqueous phase precipitated with isopropanol at −20°C for 30 min. RNA was pelleted at 14,000 × g for 20 min at 4°C, washed with 1 mL 70% ethanol, air-dried, and resuspended in 25 μL RNase-free water (Invitrogen). To remove genomic DNA, 5 μg of RNA was treated with 10 U DNase I and 50 U RiboLock RNase Inhibitor (Thermo Scientific) for 15 min at 37°C, followed by PCI (phenol–chloroform–isoamyl alcohol) extraction and ethanol precipitation with 30 mM sodium acetate (pH 5.3). Purified RNA was stored at −80°C, and quality was assessed using a Bioanalyzer RNA 6000 Nano chip (Agilent Technologies). Total RNA was also isolated from BMDMs infected with *C. albicans*, *K. pneumoniae*, both pathogens simultaneously, or non-infected controls. Co-cultures were performed at MOI 3 for 2 h, after which the medium was removed, wells were washed with PBS, and TRI Reagent LS (LabConsulting, #TS120.200) was added for RNA extraction according to the manufacturer’s instructions. For both biofilm and BMDM samples, mRNA was purified using poly-T oligo-attached magnetic beads. After fragmentation, first-strand cDNA was synthesized using random hexamer primers, followed by second-strand cDNA synthesis with either dTTP (non-strand-specific) or dUTP (strand-specific) libraries. Libraries passing quality control were sequenced as 150-bp paired-end reads on the Illumina NovaSeq 6000 platform (Novogene Sequencing Facility, UK).

### Biofilm imaging and thickness measurements

Mixed biofilms were generated as previously described by Nogueira et al with a few changes^64^. Briefly, *C.albicans* and GFP-*K. pneumoniae* were pre-culture in YPD or LB medium at 30°C and 37°C, respectively, under shaking (200 rpm), then washed twice in PBS, and 10^6^ cells were inoculated into IBIDI plates (μ-dish, 35 mm high, ibiTreat, 35 mm) containing RPMI medium (Corning, # COR50-020-PC) supplemented with 2% of glucose. Cells were allowed to adhere for 1.5 h, washed with PBS to remove non-adherent cells, supplied with fresh medium, and incubated for an additional 6 h. Prior to imaging, cells were fixed with 4% PFA for 20 min, followed by calcofluor white (CFW) staining (15 μg/mL) of single and co-cultures of *C. albicans* for 20 min in the dark, at room temperature. The fluorescence channels for CFW and EGFP were applied to image *C. albicans* and *K. pneumoniae* cells, respectively. For quantification of biofilm thickness, Z-stacks were acquired using a Zeiss LSM 980 confocal microscope equipped with an Airyscan 2 detector (8Y multiplexing mode) at the Biooptics Facility of the Max Perutz Labs Vienna. Raw Z-stacks were preprocessed by applying a mild median filter and globally normalizing intensities. To minimize local variation and account for heterogeneous biofilm structure, 100 sub-stacks (400 × 400 px) were subsampled at random positions. For each of these subsampled stacks, XZ (collapsing the y-axis) and YZ (collapsing the x-axis) projections were calculated. Each of these two projections were then segmented separately using Otsu’s method^65^ and a bounding box was fit around the segmented area. Two different thickness measurements were used: a “global” thickness value, corresponding to the height of the bounding box along the projected z-axis, and a mean thickness value, calculated from the area of segmented regions divided by the bounding-box width.

For septae estimation, stacks were maximum-intensity projected and subsequently a gaussian filter (σ=2) was applied. The single plane, projected stacks were then split into 2x2 tiles, to account for possible uneven illumination across the field of view. Images were then segmented using Yen’s method (Ref), which is generally more robust for segmentation of bright structures. Segmentation quality was visually inspected and validated to confirm accurate septa detection.

Septae labels were then counted and measured from the binary masks and counts were compared between conditions.

### Sterol and dual cell wall carbohydrate staining

For Filipin III staining, *Candida albicans* and/or *Klebsiella* spp. cells were adjusted to a final concentration of 5 × 10^6^ cells/mL and seeded at 200 µL per well onto chambered coverslips (ibidi µ-Slide, #80826), the biofilsm were let to adhere and then the media was removed and refreshed. After that, the biofilms were incubated at 37 °C for 24 h to allow maturation. Following incubation, the medium was removed and biofilms were washed three times with 200 µL PBS. Biofilms were then treated with amphotericin B (0.5 µg/mL), either alone or in combination with gentamicin (600 µg/mL), and incubated for an additional 4.5 h at 37 °C. After treatment, biofilms were washed with PBS and fixed with 100 µL of 4% paraformaldehyde (PFA) for 1 h at room temperature. Fixed samples were incubated with Filipin III (Sigma), diluted 1:20 in PBS supplemented with 10% FBS for 2 h at 37 °C. Samples were washed once with 200 µL PBS, maintained in PBS, and imaged using a Zeiss confocal microscope. For dual staining with concanavalin A (ConA) and CFW, biofilms were grown as described above. At 6 h post-adhesion, biofilms were fixed with formaldehyde and stained with Concanavalin A (25 µg/mL; Sigma, C825) for 30 min at 30 °C to visualize mannans. For concomitant analysis of chitin and mannans, samples were subsequently staining with Calcofluor White (CFW; 15 µg/mL) for 20 min in the dark. Biofilms were visualized using a Zeiss LSM confocal microscope. The Texas Red (TRed) and CFW channels were used to assess cell wall composition in single- and co-culture *C. albicans* biofilms. All images were analyzed using FIJI (ImageJ) software.

### Bioinformatics workflows

Biofilm analyses were performed as follows: paired-end reads were trimmed for adapters and low-quality bases using fastp (v0.23.2)^66^, retaining only reads with a minimum length of 20. Genomic references and coding sequences for *C. albicans* SC5314 (GCF_000182965.3) and *K. pneumoniae* subs. MGH 78578 (GCF_000016305.1) were downloaded from NCBI Datasets (https://www.ncbi.nlm.nih.gov/datasets/genome/), combined into a single FASTA file, and indexed with Salmon (v1.9.0)^67^, using a k-mer length of 31 and reference sequences as decoys. Transcript-level quantification was subsequently performed with Salmon quant, where library type was automatically inferred. All downstream analyses were conducted in the R Statistical Software (v4.2.0).^68^ Transcript-level counts were summarized to gene-level counts using tximport (v1.26.1).^69^ Genes with low expression across most of the samples were removed using the filterByExpr function and TMM normalization was applied using the calcNormFactors function from edgeR (v3.40.2).^70^ Differential gene expression analysis was carried out with limma (v3.54.2),^71^ where the mean-variance relationship was estimated and observation-level weights were computed using limma-voom prior to linear model fitting. Differentially expressed genes were identified based on empirical Bayes moderated *t*-statistics with multiple testing correction by the Benjamini-Hochberg method. Statistical significance thresholds were set at an absolute fold change >1.5 and an adjusted *p* value <0.05. Biological Process GO term enrichment was conducted using TopGO (v2.50.0),^72^ with significant terms identified by the weight01 algorithm and Fisher’s exact test. KEGG Over-Representation Analysis and Gene Set Enrichment Analysis were performed with clusterProfiler (v4.6.2).^73^ Enrichment plots were generated with gseaplot2 from the enrichplot package (v1.18.4),^74^ KEGG pathway maps were visualized with pathview (v1.38.0); Luo et al., 2013) and Venn diagrams were created using the eulerr package (v7.0.2; Larsson, 2024).

The analyses applied to the co-cultures with BMDMs were done as follows: raw paired-end reads were trimmed with *fastp* to remove adaptors, poly-N sequences, and reads with low-quality, generating the quality cutoffs: Q30, PE150, Q30 ≥ 85%, and resulting clean reads where then aligned to the mice reference genome (GRCm38/mm10) using HISAT2 (v2.0.5). Gene-level counts were generated with featureCounts (v1.5.0-p3). Differential expression analysis was performed in R (v4.1.3) with DESeq2 (v1.34.0)^75^. P values were adjusted by the Benjamini-Hochberg method to control for false discovery rate (FDR). Genes with adjusted p values < 0.05 were considered differentially expressed. Fragments per kilobase of transcript per million mapped reads (FPKM) were computed using the *DESeq2* package, based on gene length and the number of reads mapped to each gene. Functional enrichment of protein-coding DEGs (KEGG, Gene Ontology biological processes) was carried out with clusterProfiler (v4.2.2). Differentially expressed genes were visualized with ComplexHeatmap (v2.10.0)^76^ using row-wise Z-scored expression values.

### Graphics

All graphics displaying experimental setups were created using https://BioRender.com and Canva, and figures were created with Adobe Illustrator (Version 30.2.1).

### Data quantification and statistical analyses

Statistical analysis of data was performed using GraphPad Prism v10.2.3 (Graphpad Software) or R. Multi-comparison testing and kinetic comparisons was performed using One-way ANOVA followed by a Tukey post-hoc test for multiple comparisons. Weight loss curves were compared with two-way ANOVA with Geisser-Greenhouse correction and Tukey multiple comparison of each time point. Two-group comparisons were done with un-paired t test. In all cases, *p* < 0,05 was considered significant. \**p* < 0,05; \*\**p* < 0,01; \*\*\**p* < 0.001.

